# The C-terminal domain of ParB is critical for dynamic DNA binding and bridging interactions which condense the bacterial centromere

**DOI:** 10.1101/122986

**Authors:** Gemma L. M. Fisher, César L. Pastrana, Victoria A. Higman, Alan Koh, James A. Taylor, Annika Butterer, Timothy D. Craggs, Frank Sobott, Heath Murray, Matthew P. Crump, Fernando Moreno-Herrero, Mark S. Dillingham

## Abstract

The ParB protein forms DNA bridging interactions around *parS* to form networks which condense DNA and earmark the bacterial chromosome for segregation. The mechanism underlying the formation of ParB nucleoprotein complexes is unclear. We show here that the central DNA binding domain is essential for anchoring at *parS*, and that this interaction is not required for DNA condensation. Structural analysis of the C-terminal domain reveals a dimer with a lysine-rich surface that binds DNA non-specifically and is essential for DNA condensation *in vitro*. Mutation of either the dimerisation or the DNA binding interface eliminates ParB foci formation *in vivo*. Moreover, the free C-terminal domain can rapidly decondense ParB networks independently of its ability to bind DNA. Our work reveals a dual role for the C-terminal domain of ParB as both a DNA binding and bridging interface, and highlights the dynamic nature of ParB networks.

## INTRODUCTION

Bacterial chromosomes are actively segregated and condensed by the ParABS system and condensin (1). In *Bacillus subtilis*, this machinery is physically targeted to the origin proximal region of the chromosome by eight palindromic DNA sequences called *parS* (consensus sequence 5’-TGTTNCACGTGAAACA-3′) to which the ParB (Spo0J) protein binds (2). These nucleoprotein complexes act as a positional marker of the origin and earmark this region for segregation in a manner somewhat analogous to eukaryotic centromeres and their binding partners.

ParB is an unusual DNA binding protein. In addition to sequence-specific interactions with the *parS* sequence, the protein also spreads extensively around the site for about 18 kbp (2–4). The mechanistic basis for this behaviour is not well understood and a matter of active debate. Earlier models envisioned a lateral 1D spreading around *parS* to form a filament (3, 5), principally because spreading can be inhibited in a polar manner by “roadblocks” placed to the side of *parS* sequences. However, ParB foci appear to contain fewer proteins than are necessary to form a filament, and single molecule analyses using direct imaging (6) and magnetic tweezers (7) have shown that binding of DNA by ParB is accompanied by condensation. These “networks” were inferred to be dynamic and poorly-ordered, consisting of several DNA loops between distally bound ParB molecules. In cells, they are presumably anchored at *parS* sites by sequence-specific interactions but must also contain many interactions with non-specific DNA (nsDNA), as well as self-association interactions that bridge ParB protomers to form DNA loops. Modelling suggests that a combination of 1D spreading and 3D bridging interactions can explain the condensation activity and recapitulate the polar effect of roadblocks on ParB spreading (8). Recently, single-molecule imaging of the F-plasmid SopB led to a broadly similar model, defining ParB networks as fluid structures that localise around *parS* using a “nucleation and caging” mechanism (9). Despite these recent experiments converging on DNA bridging models to explain the ParB spreading phenomenon, the mechanism underpinning this behaviour remains unresolved. In particular, the relationship between these dynamic nucleoprotein complexes and the molecular architecture of the ParB protein is unclear and is the subject of the work presented here.

The genomically-encoded ParB proteins comprise three distinct domains (Figures 1A and S1A-C). Our current understanding of their structure is limited to the N-terminal domain (NTD) which binds ParA (10–13) and the central DNA binding domain (CDBD) which binds *parS* and possibly also nsDNA (14, 15). A crystal structure of *Thermus thermophilus* ParB lacking the C-terminal domain (CTD) revealed a compact dimer in which the helix-turn-helix (HtH) motifs were symmetrically arranged in a fashion that appeared suitable for binding to the palindromic *parS* sequence (Figure S1D) (14). Complementary analysis of the CTD by analytical ultracentrifugation suggested that it also formed a dimer, and it was argued that this protein interface might promote spreading interactions. More recently, a structure of *Helicobacter pylori* ParB, in which the protein was also truncated by removal of the CTD, showed a strikingly different conformation, where the NTD had moved away from the CDBD domain to form a tetrameric self-association interface (Figure S1E) (16). In this structure, the CDBD was independently bound to a *parS* half site, and it was argued that the tetramerisation of the NTD could be responsible for bridging interactions between specific and nsDNA bound to the CDBD. This has yet to be tested experimentally.

**Figure 1.**
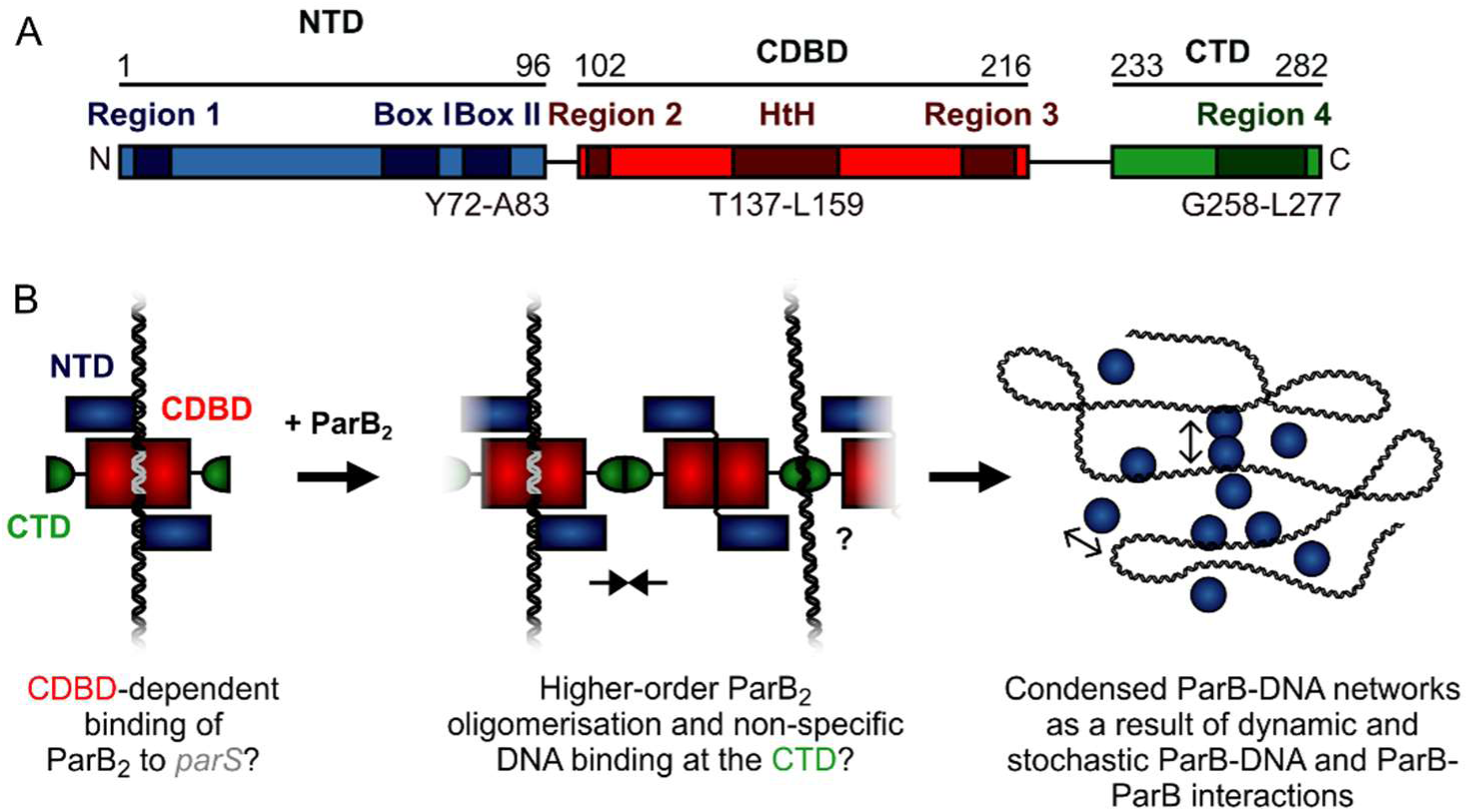
A hypothetical model for ParB-mediated condensation of the origin of replication region. (A) Domains and regions as identified in (39, 40). (B) ParB is thought to be anchored at *parS* (grey) via the HtH motif found in the CDBD (red). ParB protomers self-associate via poorly defined interactions and also make non-specific contacts with DNA segments, leading to the formation of ParB networks. In this work we have investigated the potential role of the CTD (green) in mediating ParB oligomerisation and non-specific DNA binding.

In previous work, we hypothesised that ParB contains a second DNA binding locus for nsDNA that functions independently of the helix-turn-helix motif (Figure 1B) (7). This idea was attractive to us for several reasons. Firstly, in a single DNA binding locus model, it is not straightforward to reconcile the strict localisation of ParB networks to just a few *parS* sites (and their surroundings) with the limited discrimination between specific and nsDNA binding that is observed *in vitro* (a <10-fold apparent difference in affinity) (7, 8). Secondly, although binding to *parS* protects the CDBD region from proteolysis, high concentrations of nsDNA afford no such protection, implying that it binds elsewhere on the protein (7). Thirdly, the distantly-related ParB protein from plasmid P1 provides a precedent for a second DNA binding locus in a Type I centromere binding protein, and highlights the CTD as the putative candidate region (17). However, the lack of any structural information for the CTD of a genomically-encoded ParB prevents a rigorous comparison of the systems because the primary structure similarity in this region is negligible.

In this work, we have probed the role of the CDBD and CTD of *B. subtilis* ParB using a combination of structural, biochemical, single molecule and *in vivo* approaches. We find that the CDBD is responsible for specific recognition of *parS*, and that the CTD provides both a second nsDNA binding site and a self-association interface that promotes bridging interactions and DNA condensation.

## RESULTS

### Conserved helix-turn-helix motifs in the CDBD are essential for parS recognition, but dispensable for non-specific DNA binding and condensation

Genetic and structural analyses have suggested that residue R149 may be critically important for specific binding to *parS* at the HtH locus (6, 16, 18, 19). To probe the role of the HtH motif using biochemical techniques, we compared binding of *parS* by wild type ParB and ParB^R149G^ using electrophoretic mobility shift assays containing Mg^2+^ cations (TBM-EMSA). As reported previously, inclusion of divalent cations in both the gel composition and running buffer enables the clear differentiation of specific and nsDNA-binding activities of ParB (7). As expected, binding of wild type ParB to *parS*-containing DNA produced a distinct band shift corresponding to the ParB_2_-*parS* complex, as well as poorly migrating species at high [ParB] (Figure 2A). These latter complexes are also formed on DNA that does not contain *parS*, and are therefore indicative of ParB bound to nsDNA flanking the central *parS* sequence. ParB, and mutants thereof, were purified to homogeneity (Figure S2A). EMSA experiments with ParB^R149G^ fail to produce the specific ParB2-*parS* complex whereas the formation of nsDNA complexes is largely unaffected (Figures 2A). The retention of nsDNA binding activity in ParB^R149G^ is further supported by data using gels lacking Mg^2+^ ions (TBE-EMSA) (Figure S2C), as well as a solution-based protein-induced fluorescence enhancement (PIFE) assay (Figure S2B), in which an increase in Cy3 intensity reports ParB binding. For wild type ParB, the data were fitted to the Hill equation yielding an apparent Kd of 361 ± 14 nM and Hill coefficient of 3.2 ± 0.3 in reasonable agreement with published data (7, 20). ParB^R149G^ produced a similar binding isotherm yielding a moderately weaker Kd of 493 ± 18 nM. This apparent Kd was not significantly altered when the Hill coefficient was not shared between datasets indicating the cooperativity of binding was not impaired in this ParB variant.

**Figure 2.**
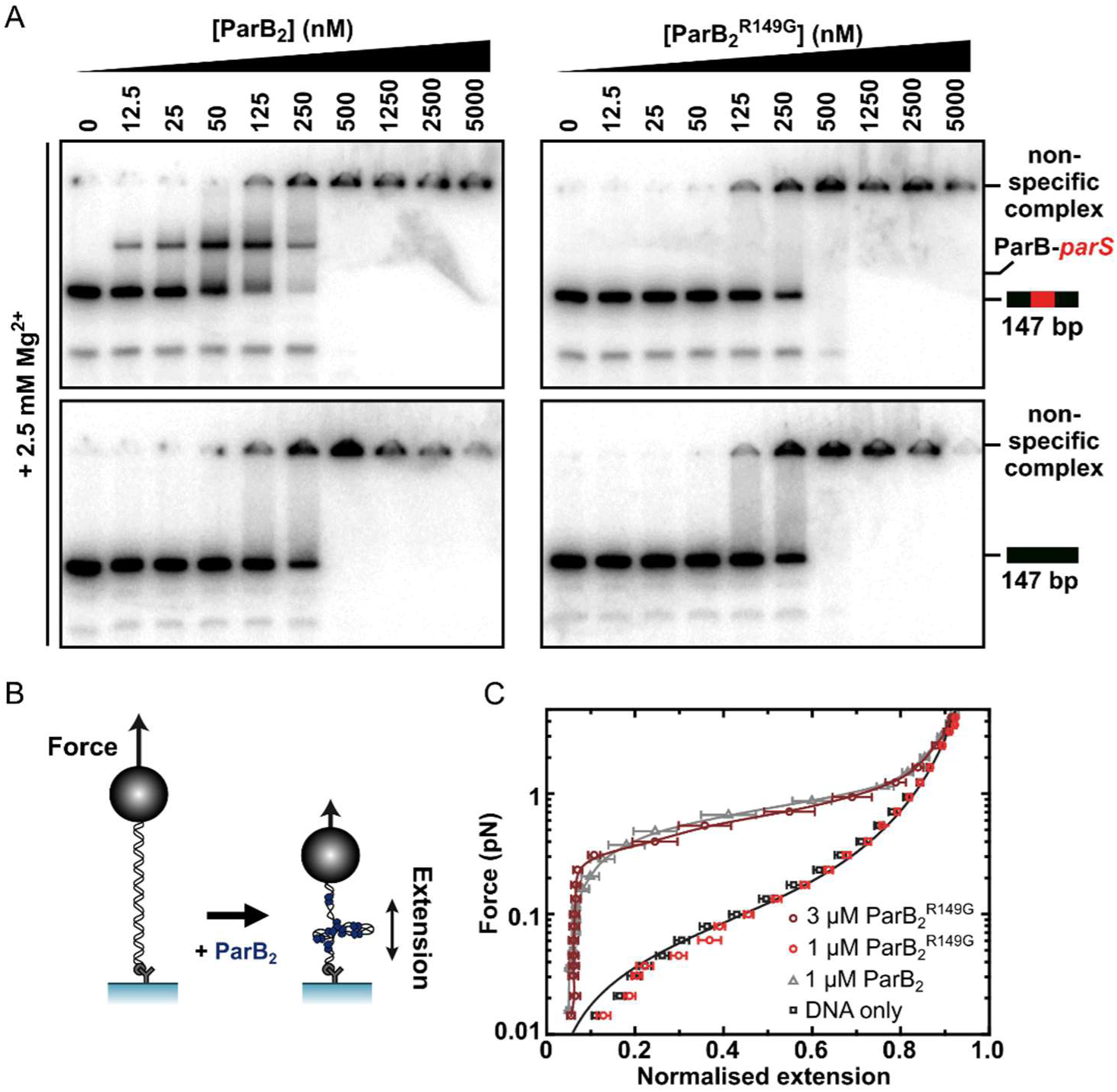
The helix-turn-helix motif is essential for specific binding to *parS*, but dispensable for nonspecific binding and condensation. (A) Representative TBM-EMSAs for wild type ParB and ParB^R149G^ monitoring binding of *parS*-containing or non-specific 147 bp dsDNA. (B) Schematic of the magnetic tweezer assay used to monitor ParB-dependent DNA condensation. (C) Mean force-extension curves for *parS*-containing DNA molecules in the presence of wild type ParB and ParB^R149G^. Non-condensed DNA data is fitted to the worm-like chain model. Solid lines in condensed data are guides for the eye. Errors are the standard error of the mean of measurements on different molecules (N∼15-35 molecules).

We next investigated the ability of ParB^R149G^ to condense DNA tethers using magnetic tweezers (Figure 2B). We previously showed that wild type ParB mediates progressive condensation of DNA substrates which is reversible by both force and protein unbinding (7). The condensed state is not highly ordered and its formation is not dependent upon *parS* sequences, indicating that nsDNA binding is sufficient for condensation. At a concentration sufficient for efficient condensation by wild type ParB (1 μM), ParB^R149G^ did not fully condense DNA, although fluctuations of the DNA tether were consistent with minor condensation events that do not greatly affect the mean extension value measured (data not shown). However, at moderately elevated concentrations (3-fold), reversible condensation did occur and was qualitatively equivalent to wild type behaviour (Figures 2C and S2D).

Together, these data show that mutation of the HtH motif effectively eliminates the ability of ParB to interact specifically with its cognate *parS* site, while nsDNA binding and condensation is only moderately affected (2-3 fold reduced). This is consistent with the idea that nsDNA binding may occur at a second DNA binding locus.

### The structure of BsParB CTD reveals a dimer with a putative DNA binding interface

We next used solution NMR to determine the structure of the ParB CTD alone (see Table S1 for structure validation and statistics and Figure S3A for an assigned ^1^H-^15^N HSQC spectrum). The structure forms a well-defined dimer containing two α-helices and two β- strands per monomer in a α1-β1-β2-α2 arrangement (Figures 3A-B). The dimer interface is formed via an intermolecular β-sheet and two domain-swapped C-terminal helices. This is somewhat similar to that seen in the P1 and SopB ParB proteins (Figures S3B-D) (15, 17), but there are also significant differences especially in the N-terminal region: the α1 helix in our structure is replaced by an additional β-strand in the CTDs of P1 ParB and SopB. Analytical ultracentrifugation, native mass spectrometry and circular dichroism thermal melt scans further confirmed that the CTD was primarily dimeric in solution and measured a Tm of 68°C (Figures S3E-F). NMR H-D exchange data revealed that the dimer is fully exchangeable (which will become important in later experiments) with the most stable H-bonds being those in the α2 helix and the intermolecular H-bonds between the two β2 strands (data not shown). These secondary structure elements are at the centre of the hydrophobic core which is made up of several Ile, Val and Phe residues in the β-sheet and several Ile/Leu residues in the α2 helix (Figure 3C). The α1 helix forms a leucine zipper with the α2 helix, where alternating Leu residues interdigitate (Figure 3D). A striking feature of the structure is a highly electropositive face of the dimer arising from several conserved Lys residues (Figures 3E-F) analogous to the plasmid-encoded SopB and P1 ParB proteins (Figure S3G-I).

**Figure 3.**
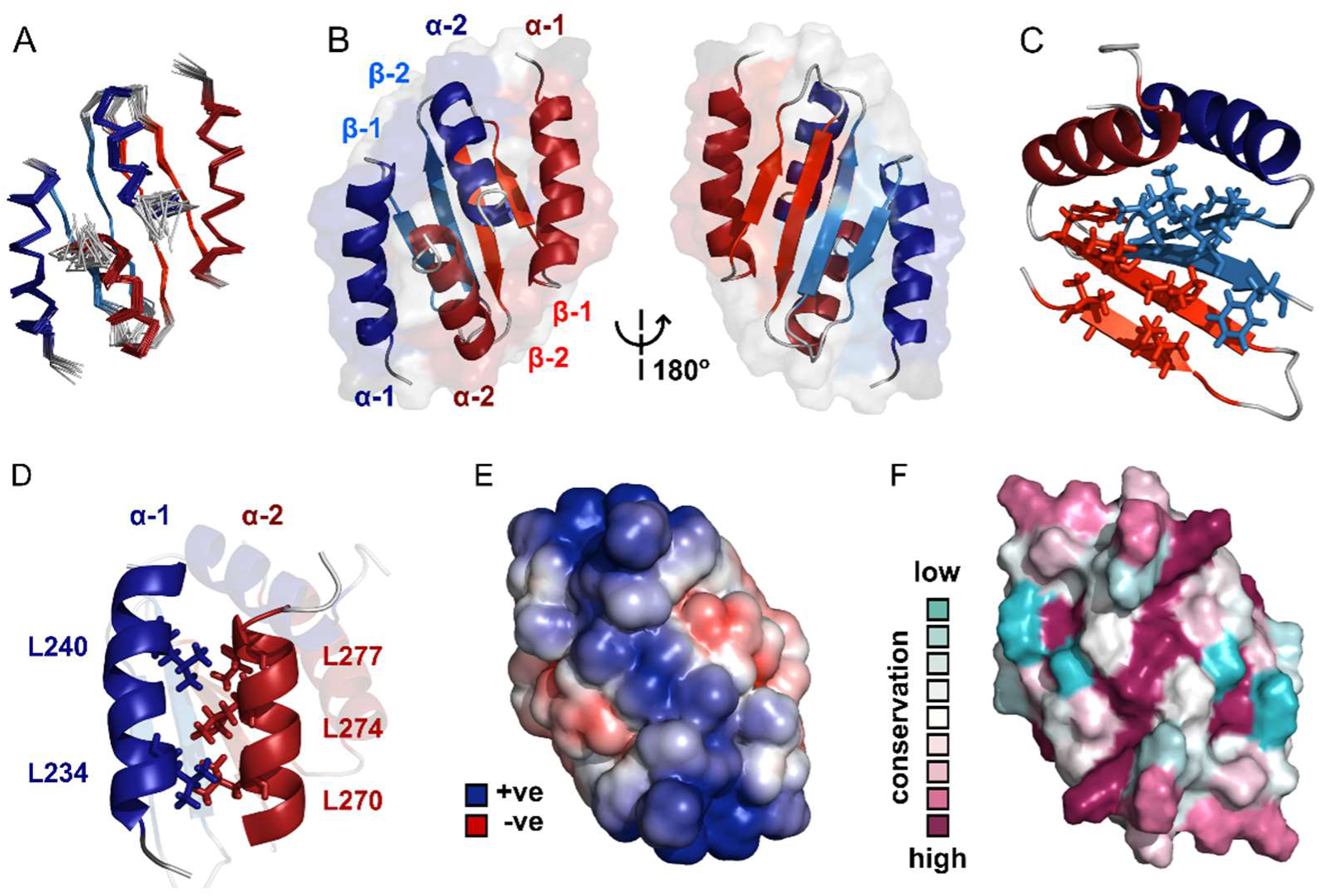
Solution NMR structure of the dimeric ParB C-terminal domain. (A) Ensemble overlay of the 14 lowest-energy CTD structures. Red and blue depict separate monomers within the dimer. (B) Secondary structure elements are identified. α1 indicates the N-terminus of each monomer. (C) The hydrophobic core of Ile, Val, Leu and Phe residues. For clarity, portions of both monomers were removed. (D) Interdigitating Leu residues of both monomers form a leucine-zipper interaction. (E) Surface charge representation reveals a large electropositive region across the β-sheet face (orientation as in B, right hand side). Continuum electrostatics calculations used the PDB2PQR web server (41) and the APBS plugin for PyMOL (42, 43). (F) Evolutionary conservation surface profile of the CTD of ParB prepared using ConSurf (44, 45) (orientation as in E). The chemical shifts, restraints and structural co-ordinates have been deposited with the BMRB and PDB for release upon publication.

### The CTD binds DNA non-specifically via a lysine-rich surface

To test the idea that the lysine-rich surface we had observed might bind to DNA, we performed TBE-EMSAs with the isolated CTD. These showed that the CTD was indeed able to bind dsDNA (Figure 4Ai) resulting in the formation of a “ladder” of bands of decreasing mobility. This is highly reminiscent of patterns formed by full length ParB under the same conditions (Figure 4Aii and (7)) except for the presence of smaller gaps between the “rungs” as would be expected for a protein of much smaller size. The CTD was also shown to bind to hairpin oligonucleotides as short as 10 bp and to ssDNA (Figure S4A and data not shown). We do not see substantial differences in the affinity of ParB for DNA substrates with different sequences and so this binding activity appears to be non-specific (data not shown). Native mass spectrometry of complexes formed between the CTD and a 15 bp duplex DNA revealed a stoichiometry of 1 DNA per dimer (Figure 4D). This is in contrast to the P1 ParB system where the CTD can bind two 16-mers (17).

**Figure 4.**
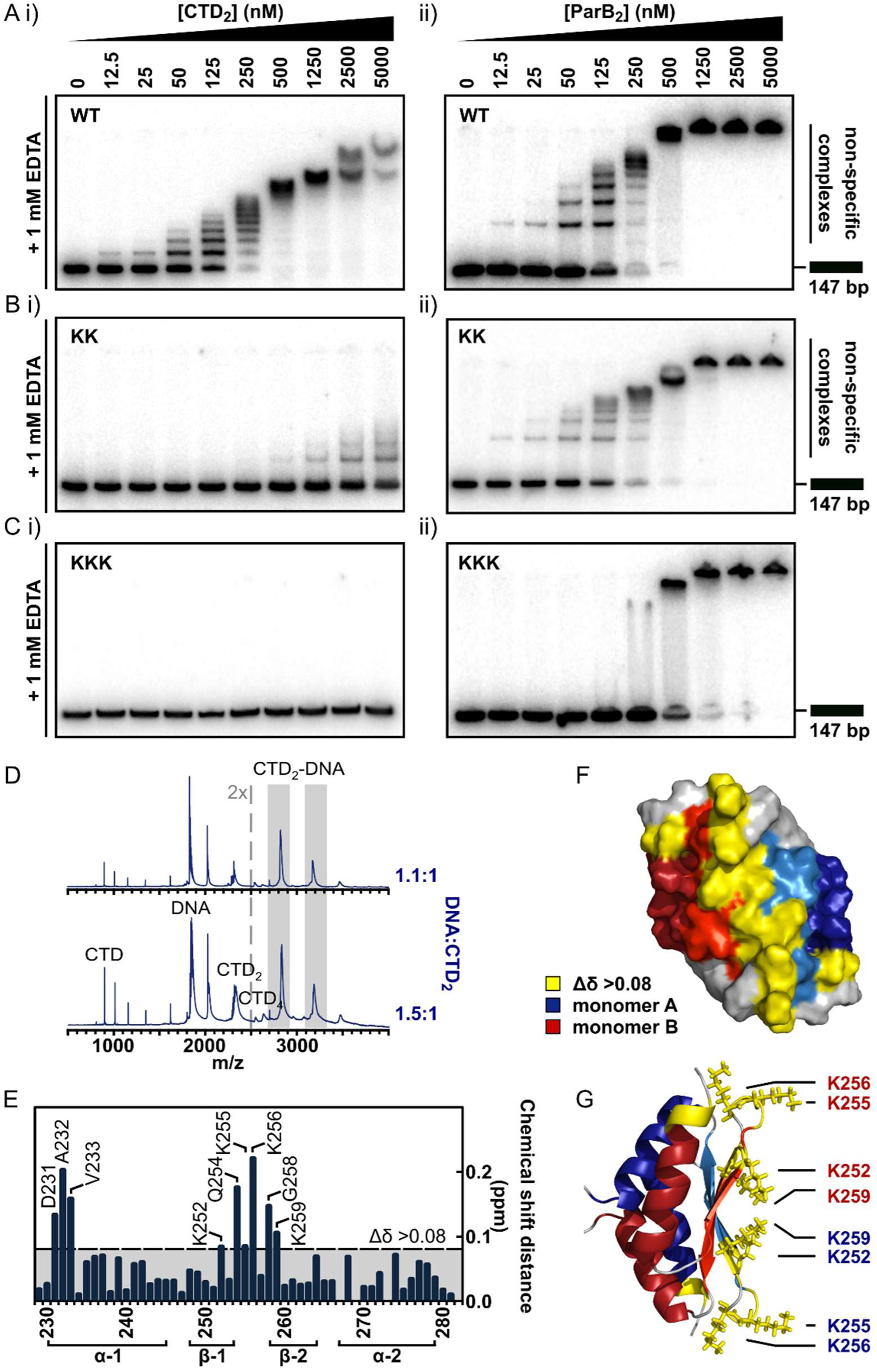
The CTD binds DNA via a lysine-rich surface. (A-C) TBE-EMSAs for the titration of full length ParB and CTD against 147 bp DNA. Wild type and mutant proteins, K255A+K257A and K252A+K255A+K259A, are indicated. (D) Native mass spectrometry. Titrations of a 15 bp nsDNA hairpin against CTD were performed between ratios of 1.1:1 and 1.5:1 DNA:CTD2. Example spectra are shown with the DNA-CTD2 complex shaded in grey. (E) Deviations in the assigned ^1^H-^15^N HSQC spectra of CTD upon titration with a 10 bp hairpin DNA. (F) Chemical shift perturbations exceeding 0.08 Δδ are highlighted on the structure in yellow. (G) Lys residues thought most likely to bind to DNA are shown as sticks.

To further probe the putative DNA binding surface, we performed a titration of the 10 bp hairpin DNA against the isotopically-labelled CTD dimer (Figure 4E, assigned ^1^H-^15^N HSQC spectra are shown in Figure S4B). Residues with large chemical shift perturbations (CSPs, Δδ >0.08) are either directly involved in DNA-binding or undergo a conformational change as an indirect result of DNA-binding, and these were mapped onto the structure (Figure 4F). Two regions of interest were identified: D231-V233, and K252-K259, which are found on the intermolecular β-sheet face and proximal loop regions to form a large, concave and positively-charged interaction surface (Figure 4G).

To confirm that this surface was responsible for DNA binding we substituted several Lys residues with Ala and monitored the effect on DNA binding using EMSA and PIFE assays. In the first instance, a dual K255A/K257A substitution was studied in the context of both the CTD-only construct (CTD^KK^) and the full length ParB protein (ParB^KK^). CTD^KK^ displayed a greatly reduced affinity (∼50-fold) for DNA, but the binding was not completely abolished (Figure 4Bi). CD thermal melt analysis confirmed that this defect was not attributable to global misfolding (Figure S4C). The full length ParB^KK^ variant showed a sigmoidal DNA binding isotherm in a PIFE assay, indicating strong positive cooperativity as observed for wild type ParB but the apparent Kd was 6-fold weaker (Figure S4D). In EMSA assays, this variant showed no defect in specific binding to *parS* as would be expected (Figure S4E). Somewhat more surprisingly however, these lysine substitutions appeared to have a negligible effect on nsDNA binding when assessed using the TBE-EMSA assays (Figure 4Bii). This may well reflect the complexity that arises when a partially defective nsDNA binding locus is physically attached to the wild type CDBD domain (which is still fully competent to bind DNA).

We next designed a triple K252A/K255A/K259A variant with the aim of fully dissipating the positive charge density across the surface of the CTD, rather than only targeting the loop-proximal regions. EMSA analysis showed that DNA binding was completely abolished in CTD^KKK^ up to concentrations of 50 μΜ (Figures 4Ci and S4F). CD thermal melt analysis showed that CTD^KKK^ was equivalently folded to wild type CTD at ambient temperatures, but with a reduced Tm (53°C) indicating a moderate destabilising effect of the mutations (Figure S4G). Interestingly, analysis of full length ParB^KKK^ showed a clear and consistent defect in all nsDNA binding assays used. TBM-EMSA gels showed that ParB-*parS* complexes were still formed (albeit in apparently lower yield, Figure S4I), whereas TBE-EMSA gels showed a complete eradication of the discrete lower mobility bands which arise from nsDNA binding (Figures 4Cii). Moreover, nsDNA binding was undetectable using the PIFE analysis (Figure S4H). Interestingly, EMSA analysis showed that DNA-bound ParB^KKK^ networks do still form as very low mobility species that form co-operatively at high ParB concentrations. This property might reflect the retention of a functional HtH motif in the ParB networks.

### DNA binding by the CTD is essential for DNA condensation and bridging *in vitro*

We next exploited our double- and triple-lysine mutant ParB proteins to test the role of the DNA-binding activity associated with the CTD in forming condensed ParB networks. These networks have been extensively characterised previously for wild type ParB using magnetic tweezers with single tethered DNA substrates (7) and also in TIRF-based microscopy (6).

Unlike full length ParB, the CTD was not capable of condensing DNA tethers under any condition tested, even up to 5 μM CTD2 concentrations and under applied forces as low as 0.02 pN (Figure S5Ai). This is consistent with the expected requirement for multiple protein-protein and/or protein-DNA interfaces to promote DNA looping and condensation. Incubation of full length ParB^KK^ with single DNA tethers resulted in defective DNA condensation compared to wild type ParB (Figure 5A). When it was observed, condensation was sudden (rather than progressive, as for wild type ParB) and full condensation required the applied force to be dropped to an exceptionally low value (0.09 pN) (Figure S5Aii). The DNA molecules also showed unusually large steps when decondensed by force, suggesting that ParB^KK^ was infrequently stabilising *in cis* DNA-bridging interactions between isolated DNA regions (data not shown). Co-incubation of ParB^KKK^ with single DNA tethers under our standard experimental conditions resulted in no measurable condensation events, even under applied forces as low as 0.02 pN and at elevated concentrations (Figures 5A and S5Aiii). The average work done by ParB compared to the variant proteins during these condensation events was determined from the difference between the integral of the force-extension curve in the presence of the protein and that of DNA alone. This provides a means to quantitatively compare the condensation efficiency between mutants (Figure S5B). We also performed plectoneme stabilisation experiments (Figure 5B). In this assay, a single torsionally-constrained DNA molecule is positively supercoiled at a 4 pN force by applying 60 turns. ParB is then introduced and, after full buffer exchange, all turns are released whilst monitoring DNA extension. Any deviation of DNA extension from that expected of bare DNA is indicative of supercoiled regions being stabilised by ParB. ParB^KK^ could stabilise DNA-bridging interactions between isolated DNA regions but this was often characterised by large steps in the DNA tether extension increase which is unlike the behaviour of wild type (Figures 5C and S5C). ParB^KKK^ was unable to stabilise plectoneme structures showing that it cannot bridge DNA segments *in trans* (Figures 5D and S5D).

**Figure 5.**
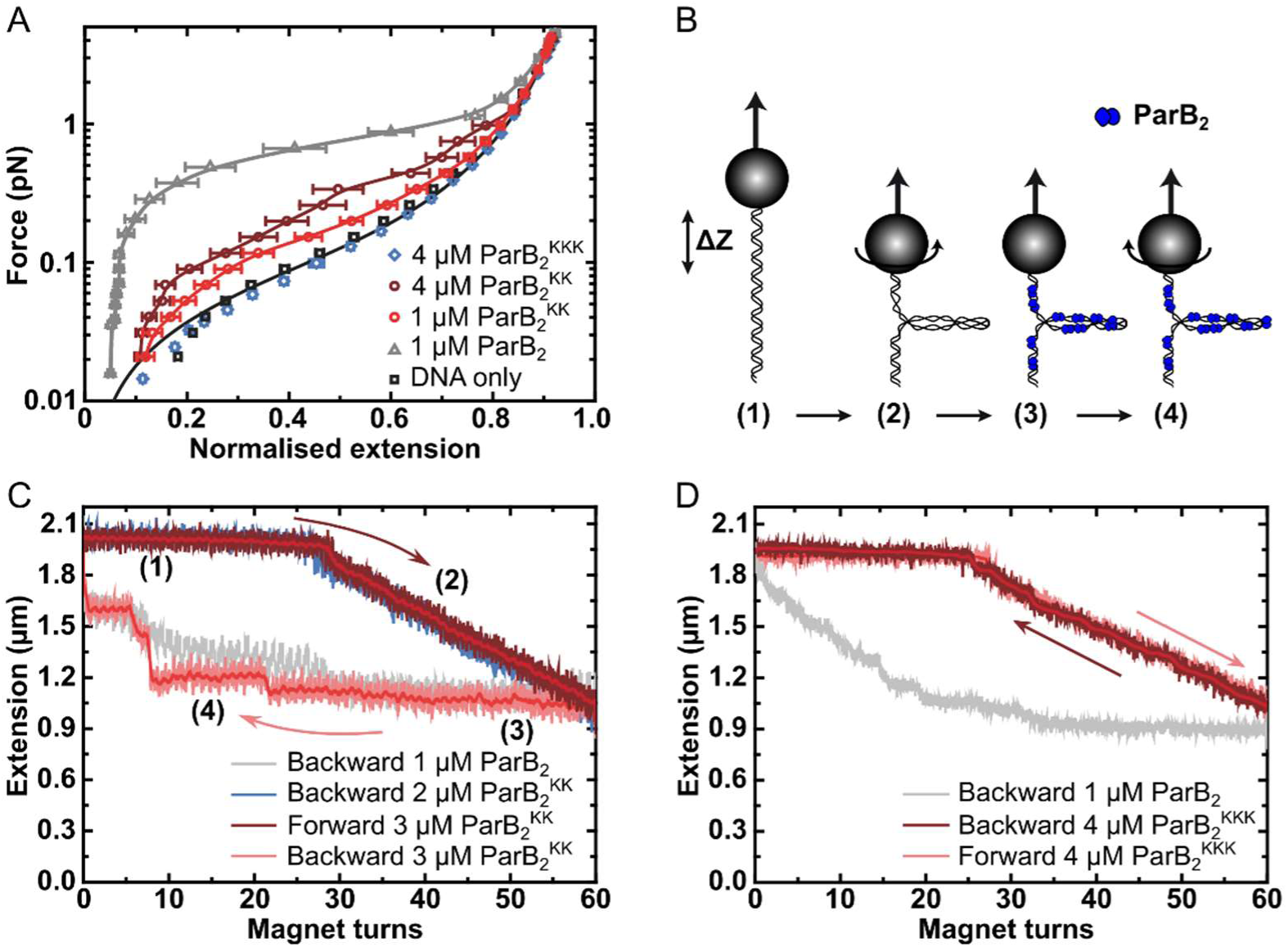
DNA binding by the CTD is required for efficient DNA condensation *in vitro*. (A) Mean force-extension curves of DNA molecules co-incubated with ParB variants at the indicated concentrations. Non-condensed (protein-free) DNA data is fitted to the worm-like chain model. Solid lines in condensed data are guides for the eye. Errors are the standard error of the mean of measurements on different molecules (N∼18-35 molecules). (B) Schematic of plectoneme stabilisation assay. A single torsionally-constrained DNA molecule was positively supercoiled at 4 pN force by applying 60 turns. This shortens the tether length due to the formation of plectonemes in the overwound DNA. ParB**2** is then introduced and all turns are released whilst monitoring DNA extension. Evidence for ParB-dependent plectoneme stabilisation is provided by hysteresis in the extension as a function of magnet turns as the supercoiling is removed. (C) Plectoneme stabilisation assay comparing bare DNA, wild type ParB and ParB^KK^. The double-mutant protein supported DNA bridging and occasionally large steps were observed in the backward trace (see text for discussion). D) Plectoneme stabilisation assay comparing wild type ParB and ParB^KKK^. No activity was detected for the triple-mutant protein.

### The CTD can both inhibit the formation of, and decondense, ParB-DNA networks *in vitro*

The CTD potentially acts as both an oligomerisation interface and also a site of nsDNA-binding. Therefore, we hypothesised that the CTD might have a dominant negative effect on full length ParB by competing for the DNA and protein interfaces that mediate the formation of ParB networks in the magnetic tweezers.

Purified CTD completely inhibited the formation of the condensed state if pre-incubated with wild type ParB and DNA under the high stretching force regime (Figures 6A and S6A). We also tested whether the introduction of free CTD to pre-condensed tethers was able to disrupt ParB-DNA networks. Condensed ParB networks were completely stable in a flow of free ParB on the timescale of these experiments, and the DNA tethers were also able to recondense following force-induced decondensation (Figure 6Bi). However, the inclusion of excess free CTD rapidly disrupted ParB networks, with some degree of decondensation observed in 94% of all the molecules tested (Figure 6Bii). Moreover, those molecules which did not decondense spontaneously could be stretched by force, but were then unable to recondense when permissive forces were restored. This ability of the CTD to decondense ParB networks demonstrates that the protein-protein and/or protein:DNA interfaces that maintain the condensed state under a low force regime are dynamic (i.e. they are exchanging while the overall structure of the network is maintained).

**Figure 6.**
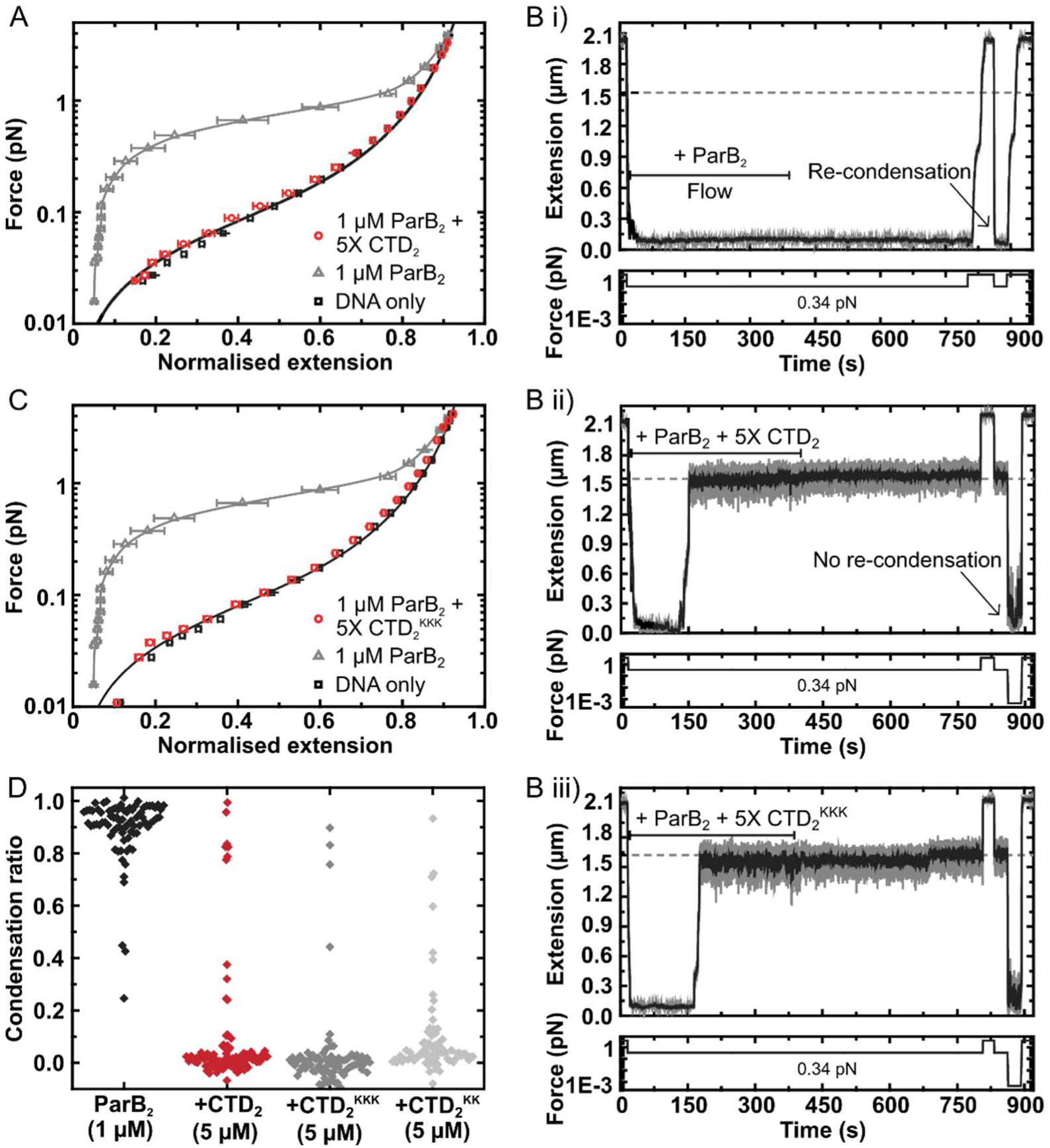
The CTD of ParB both inhibits and disrupts ParB-dependent DNA condensation. (A) Mean force-extension curves for DNA molecules co-incubated with 1 μM ParB2 in the presence or absence of 5 μM CTD2. Non-condensed (protein-free) DNA data is fitted to the worm-like chain model. Solid lines in condensed data are guides for the eye. (B) i) ParB-DNA networks are stable in magnetic tweezers in the presence of 1 μM ParB2. ii) ParB-DNA complexes spontaneously decondense following the introduction of 1 μM ParB2 and 5 μM free CTD2. iii) ParB-DNA complexes spontaneously decondense following the introduction of 1 μM ParB2 and 5 μM CTD2^KKK^. (C) Mean force-extension curves for DNA molecules co-incubated with 1 μM ParB2 in the presence or absence of 5 μM mutant CTD2^KKK^. Errors are the standard error of the mean of measurements on different molecules (N>40 molecules) (D) Condensation ratio (see Methods for definition) for individual DNA condensation events involving the addition of CTD competitor variants to pre-condensed ParB-DNA networks.

We have shown above that the CTD binds tightly to nsDNA. Therefore, its ability to prevent condensation and induce decondensation might simply reflect competition for the nsDNA that becomes available during exchange of ParB:DNA interfaces. Indeed, we have shown previously that free DNA is a potent inducer of network decondensation in the MT apparatus (7). To test the idea that the CTD dimerisation interface is also important for maintaining the condensed state, we repeated our experiments with the CTD^KK^ and CTD^KKK^ constructs, which are defective and apparently unable (respectively) to bind nsDNA. Both mutant proteins were as effective as wild type in preventing condensation (Figures 6C and S6B), and both were able to induce decondensation in approximately 95% of all molecules tested (Figures 6Bii and 6D). This strongly suggests that CTD-dependent ParB network dissipation is primarily mediated by competition for the CTD dimerisation interface and further confirms that the CTD^KKK^ construct is folded. This competition presumably results from the formation of heteroligomers between full length ParB and the CTD, which disrupts interactions that are essential for condensation (Figure S6C).

### The CTD is critical for the formation of ParB foci *in vivo*

To test the importance of the CTD dimerisation and DNA binding interfaces in *vivo*, we compared the ability of wild type and mutant ParB-GFP proteins to form foci in *B. subtilis* cells when expressed from the endogenous locus. Wild type ParB-GFP formed discrete foci around *oriC* as expected (3, 6, 18, 21–25). In contrast, ParB^KKK^-GFP failed to form discrete foci (Figure 7Ai-ii) despite wild type expression (Figure S7A). Interestingly, the triple-mutant protein appeared to localise non-specifically to the nucleoid, perhaps as a result of residual DNA binding by the HtH motifs, suggesting that ParB^KKK^-GFP retained the ability to dimerise. A parB^L270D+L274D^ construct, designed to prevent dimerisation of the CTD, was completely unable to form ParB foci (Figure 7Aiii-v) despite being expressed at approximately wild type levels (Figure S7B). The complete deletion of the CTD by truncation to E222 or E227 resulted in the same phenotype (data not shown).

**Figure 7.**
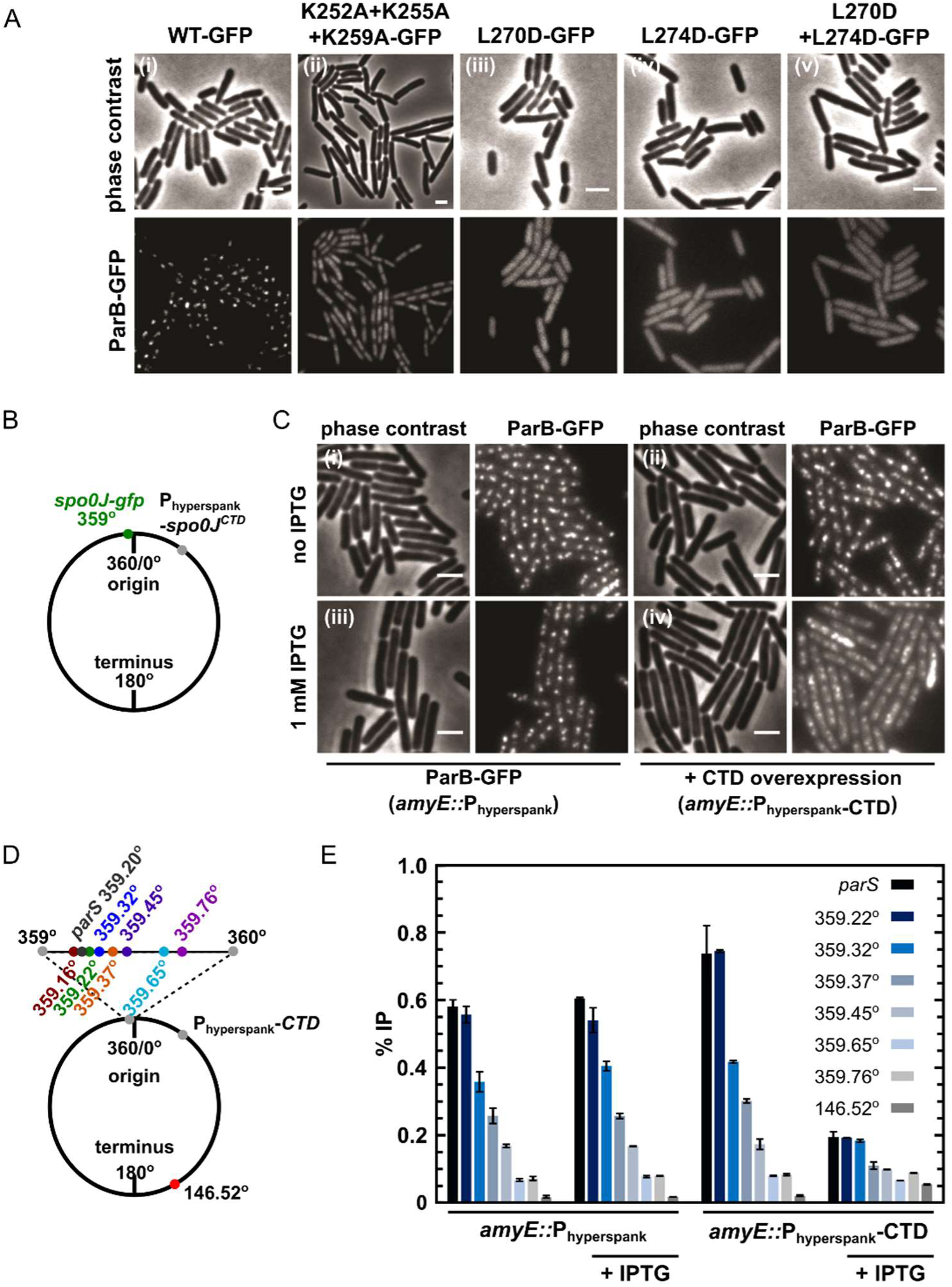
DNA-binding and dimerisation by the CTD is critical for ParB function *in vivo*. (A) ParB-GFP mutants form abnormal foci in *B. subtilis*. Cells were grown overnight in slow growth conditions before dilution (1:100) into fast growth media, and were allowed to achieve at least 5 mass doublings before observation by microscopy. (B) Construct design for overexpression of the CTD *in vivo*. (C) CTD overexpression was induced by IPTG in the presence of chromosomally-encoded wild type ParB-GFP using the Phyperspank construct. Cells were grown as in A. (D) Construct design for ChIP-qPCR. (E) ChIP-qPCR assay for ParB spreading. Cells were grown slowly overnight, diluted (1:100) into fast growth media, and allowed to reach 8 mass doublings before crosslinking with 1% formaldehyde. Background IP was measured at the terminus (146°). Primer pairs produced 200-300 bp fragments.

Our attempts to purify recombinant parB^L270D+L274D^ failed because the protein was insoluble upon overexpression in *E. coli*. This raises the possible caveat that the loss of function associated with this dimerisation mutant *in vivo* might reflect mis-folding. Therefore, we also investigated whether the free CTD was able to interfere with dimerisation in *vivo*, thereby causing a dominant negative effect on ParB function. A *B. subtilis* strain was engineered with a C-terminal *gfp* fusion replacing the endogenous *spo0J* and the unlabelled CTD-only gene inserted at an ectopic locus downstream of a Phyperspank promoter, designed for high protein expression that is tightly-controlled with IPTG (Figure 7B). Overexpression of the CTD caused ParB-GFP foci to become diffuse (Figure 7C), although expression levels of endogenous ParB-GFP were unaffected (Figure S7C).

ChIP-qPCR analysis allowed us to more directly characterise the effect of CTD expression upon ParB spreading (i.e. the enrichment of ParB at and widely around *parS* sites). Spreading was measured around a single *parS* site (359.20°) and used a locus towards the terminus to monitor background ‘enrichment’ (146.52°) (Figure 7D). As expected, in the absence of CTD expression, ParB was highly enriched not only at *parS* sites (∼40-fold), but also for several kilobase pairs around *parS* (Figure 7E). Overexpression of CTD significantly decreased the signal around *parS* (∼4-fold enrichment), indicating that it interferes with spreading. Western blotting of cells grown under equivalent conditions to the ChIP-qPCR assay and using the same batch of polyclonal anti-ParB antibody suggests that CTD is not preferentially recognised over the endogenous ParB copy (Figure S7D). Note that the reduced signal observed for the *parS* fragment does not necessarily indicate defective specific binding because the PCR product at *parS* is much larger than the 16 bp *parS* site or the 24 bp footprint of a ParB dimer (3).

Finally, we determined the consequence of decreased ParB spreading *in vivo* induced by CTD overexpression by measuring the rate of DNA replication initiation. ParB normally inhibits the activity of Soj, a regulator of the master bacterial initiation protein DnaA (26). Marker frequency analysis showed that CTD overexpression stimulated the frequency of DNA replication initiation, indicating that regulation of DnaA by Soj was adversely affected (Figure S7E). Together, these results are consistent with our *in vitro* observations, and support a model in which dynamic ParB-DNA networks are dependent upon ParB oligomerisation and DNA-binding interfaces in the CTD.

## DISCUSSION

ParB proteins form long-distance bridging interactions on DNA substrates to form foci that facilitate chromosomal partitioning reactions (4, 5, 27). These ParB foci are anchored at *parS* sites and interact non-specifically around a single site for ± 18 kbp (2, 28). This ParB “spreading” activity appears to be a conserved property across chromosomal and plasmid segrosomes, yet the protein interaction interfaces involved have remained elusive, particularly for the genomically-encoded systems (6). This is, in part, due to the highly variable structures of ParB proteins and their cognate centromere sequences, even within the type I subclass of which *B. subtilis* ParB is a member (29, 30). Increasing evidence indicates that ParB spreading is the result of a DNA-bridging activity mediated by ParB-ParB oligomerisation interfaces (6–9, 31). However, a complete understanding of the relationship between ParB structure and function has been hindered by the lack of any full length structure for a chromosomally-encoded ParB. Indeed, the organisation of the N-terminal (NTD), central DNA-binding (CDBD) and C-terminal (CTD) domains appears to be quite complex (Figures S1A-C). For the type I ParB protein class, there is evidence to suggest that dimerisation and/or tetramerisation can occur at the NTD and CDBD, and that dimerisation can occur at the CTD (14–16). A combination of some or all of these activities must support ParB oligomerisation.

To directly address the putative role of the CTD in spreading (14), we resolved the first structure of a genomic ParB CTD. The structure revealed a conserved lysine-rich surface and we showed that this binds to DNA in an apparently non-specific manner. This novel DNA binding locus is structurally distinct from the sequence-specific DNA binding site for *parS* formed by the classical helix-turn-helix motif within the CDBD domain. In this respect, there are parallels with the plasmid-encoded ParB proteins P1 ParB and SopB (17, 32). In all of these systems, the CTD shares similar surface electrostatics in which a polar distribution of charged residues results in both positively- and negatively-charged surfaces on opposite faces of the domain. In the case of both *B. subtilis* ParB and P1 ParB, the Lys/Arg rich surface has been shown to bind directly to DNA using structural and/or biochemical techniques ((15, 17) and this work). The integrity of the CTD may also be important for stabilising the N-terminal region of the protein. When SopB was truncated ahead of the C-terminal domain boundary, it could not bind *sopC* (the F-plasmid equivalent of *parS*) (33), and analogous results have been obtained with *T. thermophilus* and *B. subtilis* ParB ((14) and unpublished observations).

Our CTD structure facilitated the design of separation of function mutations to test the importance of the dimerisation and DNA binding activities using a variety of *in vitro* and *in vivo* readouts of ParB function. We showed that the CTD is not required for *parS* binding, and that this is instead dependent on the HtH motif found within the CDBD domain as predicted in several previous studies (6, 16, 18, 19, 29, 34–36). In contrast, the CTD is essential for the formation of nsDNA complexes that are observed as ladders of decreasing mobility in EMSA assays. Mutant proteins that were unable to bind DNA at the CTD locus were severely defective in both DNA condensation assays *in vitro* and ParB foci formation assays *in vivo*. Moreover, ParB proteins that were designed to be unable to form oligomeric structures by mutation of the CTD-CTD dimerisation interface were completely unable to form ParB foci and the free CTD domain was able to disrupt ParB networks independently of its ability to bind DNA. Together, our observations strongly support the idea that the CTD is essential for both DNA binding and for ParB-ParB bridging interactions that support DNA condensation.

We propose that the presence of two DNA binding loci in ParB can help to explain how ParB networks are anchored at *parS in vivo*. Importantly, this architecture resolves the paradoxical observation that the apparent specificity for *parS in vitro* (<10-fold greater affinity for *parS* versus nsDNA) is insufficient to explain the strict localisation of ParB around just 8 sites in a ∼4 Mbp genome (7, 8). In a two DNA binding site model, specific and non-specific binding can be semi-independent activities that are architecturally-coupled only when ParB oligomerises into networks. This model can also explain why DNA condensation does not require *parS in vitro*, whereas the absence of *parS* sites prevents the formation of ParB-DNA foci *in vivo* ((6, 7, 9, 37) and this work). In a test tube, whenever ParB is present at concentrations that licence oligomerisation, it is always in large stoichiometric excess over binding sites and all available DNA will be bound. In cells, the situation is very different because there is a limited pool of ParB (6). Specific interaction with *parS* preferentially anchors the ParB network at *parS*, leaving a vast number of unoccupied sites. If *parS* sites are absent in cells, ParB might still form networks, but these would not be anchored at specific sites and would therefore fail to form foci, as has been observed experimentally (37, 38). A rigorous proof of these ideas will require a modelling approach that will be the subject of future work.

Previously, high-resolution SIM and ChIP-seq data have suggested that ParB-DNA partition complexes involve stochastic and dynamic binding of ParB to both DNA and other ParB proteins, resulting in the formation of fluid intra-nucleoid “ParB cages” on DNA (9). This view is consistent with the disorder observed in magnetic tweezers assays (7), and with the dominant negative effect of the free CTD domain on ParB networks shown here. However, a recent structural study of *H. pylori* ParB concluded that a novel tetramerisation interface within the NTD was also likely to be important in bridging (16). Moreover, spreading could be facilitated by *parS*-dependent conformational changes that act as nucleation points for networks (8, 14). A more complete understanding of ParB network formation and its regulation will be required to underpin future studies on how ParB acts together with ParA and condensin to orchestrate efficient chromosome segregation.

## ACKNOWLEDGMENTS

We thank Benjamin Bardiaux for providing the latest version of the Aria software which included the softened force field, and also Chris Williams, Joe Beesley and Antony Burton for assistance with NMR, CD and AUC data acquisition, respectively. G.L.M.F. was supported by a PhD studentship from the Biotechnology and Biological Sciences Research Council (grant number 1363883). M.S.D., J.A.T., V.A.H. and T.D.C. were supported by the Wellcome Trust (grant numbers 100401 and 077368). F.M.-H. acknowledges support from the European Research Council (ERC) under the European Union’s Horizon 2020 research and innovation (grant agreement No 681299) and from Spanish MINECO (ref. FIS2014-58328-P).

## AUTHOR CONTRIBUTIONS

G.L.M.F. C.L.P., F.M.-H. and M.S.D. designed the study. G.L.M.F. carried out all protein preparation and biochemical analysis under the supervision of M.S.D. and T.D.C. C.L.P. performed magnetic tweezers experiments under the supervision of F.M.-H. V.A.H. resolved the structure using NMR under the supervision of M.P.C. A.K. carried out all *in vivo* analysis under the supervision of H.M. J.A.T. carried out preliminary work on ParB^R149G^ A.B. undertook native mass spectroscopy under the supervision of F.S. Manuscript was written by G.L.M.F. and M.S.D. with input from all authors.

## SUPPLEMENTARY INFORMATION

### METHOD DETAILS

#### Plasmids and DNA substrate preparation

All mutagenesis used the pET28a-ParB expression vector as a template (7). The R149G mutation was introduced by site-directed mutagenesis using a QuikChange II XL kit (Agilent Technologies). The full length ParB gene (1-849) with the K255A+K257A or K252A+K255A+K259A substitutions was produced synthetically (Life Technologies) and subcloned into pET28a using NcoI and BamHI restriction sites (7). CTD only (217-282) constructs were produced using PCR with primer overhangs incorporating 5’ PacI and 3’ XmaI restriction sites for subcloning (5’ - GCGTAAGCCCCGGGCAGAATGTTCCACGTGAAACAAAG – 3’ and 5’ - GCGTCATGTTAATTAATCATTATGATTCTCGTTCAGACAAAAG – 3’) into pET47b (Novagen) to produce a protein with an N-terminal HCV 3C protease cleavable His-tag. The integrity of all DNA sequences was confirmed by direct sequencing (DNA Sequencing Service, University of Dundee).

Preparation of radiolabelled, 5’Cy3-labelled and magnetic tweezer DNA substrates was as described (7). 10 bp DNA hairpins were prepared by heating a self-complementary oligonucleotide (5′ - GCGTACATCATTCCCTGATGTACGC - 3′) in 10 mM Na_2_HPO_4_/NaH_2_PO_4_ pH 6.5, 250 mM KCl and 5 mM EDTA to 95 °C for 25 mins, followed by rapid cooling in an ice bath. The DNA was purified by anion-exchange chromatography using a 0.25-1 M KCl gradient, and desalted over multiple NAP-10 columns (GE Healthcare Life Sciences) before concentration in a centrifugal vacuum concentrator.

#### ParB overexpression and purification

ParB, and the variants R149G, K255A+K257A and K252A+K255A+K259A, were overexpressed and purified as described (7). CTD, and mutants thereof, were His-tagged and purified to homogeneity as follows. Cell pellets, produced as described (7), were resuspended in 20 mM Tris-HCl pH 7.5, 500 mM NaCl and 1 mM BME (TNB buffer) with the addition of 10 mM imidazole, 5% (v/v) glycerol and protease inhibitor cocktail set II (Millipore), before being snap-frozen and stored at -80 °C. Cells were lysed by sonication in the presence of 0.2 mg/ml lysozyme (Sigma). The lysate was clarified by centrifugation and loaded on to a 5 ml HisTrap HP column (GE Healthcare Life Sciences) equilibrated with TNB buffer + 10 mM imidazole. CTD elution was achieved with a linear gradient of 10 mM to 500 mM imidazole. Peak fractions were assessed by SDS-PAGE and pooled accordingly. The tag was removed with HRV 3C protease (Thermo Scientific, Pierce) for 16 hrs at 4 °C during dialysis into TNB buffer + 10 mM imidazole. The products were subsequently loaded onto a HisTrap HP column whereby the cleaved CTD was collected in the flow-through volume, followed by concentration by centrifugation in Amicon Ultra-15 3 kDa MWCO spin filters (Millipore). This concentrate was loaded at 1 ml/min onto a Hiload 16/600 Superdex S75 gel filtration column (GE Healthcare Life Sciences) equilibrated with 50 mM Tris-HCl pH 7.5, 1 mM EDTA, 300 mM NaCl and 1 mM DTT (storage buffer). Appropriate peak fractions were pooled, followed by final concentration by centrifugation as described. Spectrophotometric grade glycerol (Alfa Aesar) was added to 10% (v/v). The final protein was then snap-frozen as aliquots and stored at -80 °C. ParB concentration was determined by spectrophotometry using theoretical extinction coefficients of 7450 M^-1^ cm^-1^ and 2560 M^-1^ cm^-1^ for ParB and CTD respectively. ParB concentrations in all assays refer to the dimeric state.

For structure determination by NMR, the CTD was dual isotopically (13C and 15N) labelled during overexpression in M9 media, as described previously (46), and subsequently purified as above.

#### CD spectroscopy

CD spectra were collected using a JASCO J-810 spectropolarimeter fitted with a Peltier temperature control (Jasco UK). 50 μM protein samples were buffer exchanged into phosphate buffered saline (PBS; 8.2 mM NaH_2_PO_4_, 1.8 mM KH_2_PO_4_, 137 mM NaCl and 2.7 mM KCl (pH 7.4)) by 16 hr dialysis at 4 °C using a membrane with a MWCO of 3.5 kDa. At 20 μM and using a 0.1 cm quartz cuvette, thermal stability data was acquired across a 190-260 nm absorbance scan (1 nm data pitch at a scanning speed of 100 nm/min) from 5 to 90 °C at 5 °C increments. Raw data was normalised to molar ellipticity (MRE (deg.cm^2^.dmol^-1^)) using calculation of the concentration of peptide bonds and the cell path length. A buffer only baseline was subtracted from all datasets.

#### NMR

NMR datasets were collected at 35 °C, utilising a Varian VNMRS 600 MHz spectrometer with a cryogenic cold-probe. The purified protein was buffer exchanged into PBS (10 mM NaH_2_PO_4_, 1.8 mM KH_2_PO_4_, 137 mM NaCl and 2.7 mM KCl (pH 6.1)) and concentrated to 1 mM. ^1^H-^15^N HSQC, ^1^H-^13^C HSQC, HNCACB, CBCA(CO)NH, HNCA, HN(CO)CA, HNCO, CC(CO)NH, H(CCO)NH, HCCH-TOCSY, ^15^N-NOESY-HSQC (150 ms mixing time), ^13^C-NOESY-HSQC (140 ms mixing time) and aromatic ^13^C-NOESY-HSQC (140 ms mixing time) experiments were collected on ^13^C,^15^N-labelled CTD. 2D ^1^H-^1^H TOCSY and NOESY spectra were recorded on unlabelled protein. ^13^C,^15^N-labelled and unlabelled protein were mixed in equimolar amounts to create a mixed labelled sample used to record 3D ^13^C, ^15^N F_1_-Filtered, ^13^C, ^15^N F3-edited ^13^C-NOSEY-HSQC and ^15^N-NOESY-HSQC experiments (47). A hydrogen-deuterium (HD) exchange experiment was conducted by recording ^1^H-^15^N HSQC experiments at several intervals following dissolution of freeze-dried protein in D_2_O. A titration was conducted by adding a 10 bp DNA hairpin step-wise to ^13^C, ^15^N-labelled CTD and recording a ^1^H-^15^N HSQC experiment after each addition. The final molar ratio of protein:DNA was 1:1.25. All NMR data were processed using NMRPipe (48). Spectra were assigned using CcpNmr Analysis 2.4 (49). Proton chemical shifts were referenced with respect to the water signal relative to DSS.

Heteronuclear NOE experiments showed residues 214-228 to be highly flexible. This was supported by chemical shift analysis with TALOS+ (50) and the absence of any medium or long-range NOEs for these residues. Structure calculations were only conducted on residues 229-280, as the unstructured tail made unfavourable energy contributions to the calculation which distorted the selection of ensembles of low-energy structures. Structure calculations were conducted using ARIA 2.3 (51). 10 structures were calculated at each iteration except iteration 8, at which 200 structures were calculated. The 20 lowest energy structures from this iteration went on to be water refined. Spin diffusion correction was used during all iterations (52). Two cooling phases, each with 30,000 steps were used. Torsion angle restraints were calculated using TALOS+. Standard ARIA symmetry restraints for 2 monomers with C2 symmetry were included (53). Structural rules were enabled, using the secondary structure predictions made by TALOS+. The HD exchange experiment showed 29 NH groups to be protected after 8 mins. Initial structure calculations were conducted without hydrogen bond restraints. Hydrogen bond donors were then identified and corresponding hydrogen bond restraints included in later calculations. Calculations were conducted using a flat-bottom harmonic wall energy potential for the distance restraints until no consistent violations above 0.1 Å were observed. The final calculation was then performed using a log-harmonic potential (54) with a softened force-field (55). Structures were validated using the Protein Structure Validation Software (PSVS) suite 1.5 (56) and CING (57). The chemical shifts, restraints and structural co-ordinates have been deposited with the BMRB and PDB for release upon publication.

#### EMSA experiments

The specific and nsDNA-binding activity of ParB was analysed by TBM- and TBE-PAGE as described (7). Serial dilutions of ParB, to the indicated concentrations, were incubated with 20 nM 147 bp *parS* or “scrambled” DNA (at a ratio of 1:19 labelled to unlabelled), 50 nM HEPES-KOH pH 7.5, 100 mM KCl, 2.5 mM MgCl_2_, 0.1 mg/ml BSA, 1 mM DTT and 2.5 % (v/v) Ficoll in a 20 μl reaction volume. Where indicated, different length dsDNA substrates were used equivalently. Samples were incubated at room temperature for 30 mins followed by 5 mins on ice. 10 μl of each were loaded onto a 6% acrylamide/bis-acrylamide (29:1) gel in 90 mM Tris, 150 mM H_3_BO_3_ (final pH 7.5), supplemented with either 2.5 mM MgCl_2_ (TBM) or 1 mM EDTA (TBE). Gels were pre-run at 150 V, 4 °C for 30 mins in a buffer identical to their composition, and run post-loading at 150 V, 4 °C for 1 hr. For imaging, gels were vacuum-dried, exposed to a phosphor screen and subsequently scanned by a Phosphor-Imager (Typhoon FLA 9500, GE Healthcare Life Sciences).

#### Protein Induced Fluorescence Enhancement (PIFE) assay

ParB DNA-binding to non-specific substrates was analysed in a solution-based assay where a change of emitted Cy3 fluorescence acted as a reporter of ParB binding (7). ParB was incubated with 20 nM 147 bp 5’-Cy3-labelled DNA, 50 mM HEPES-KOH pH 7.5, 100 mM KCl, 2.5 mM MgCl_2_, 0.1 mg/ml BSA and 1 mM DTT. Samples of 120 μl were incubated at room temperature for 30 mins before being transferred into a quartz cuvette for data collection. Cy3 fluorescence in each sample was measured by excitation at 549 nm and an emission scan between 560 and 600 nm (Cary Eclipse Fluorescence Spectrophotometer, Agilent Technologies). Peak maxima were calculated by the area under the curve function in GraphPad Prism software, and the increase in fluorescence calculated relative to a DNA only control. Where appropriate, data was fitted with a Hill equation.

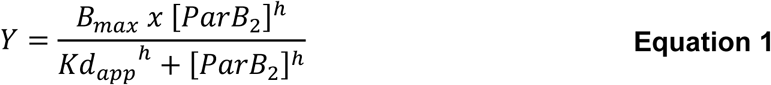
Where Y is the measured increase in fluorescence, B_max_ is the maximal increase in fluorescence, h is the Hill coefficient and Kd*_app_* is the apparent dissociation constant. When comparing wild type and mutant binding isotherms, the data were well-fitted using a shared value for the Hill coefficient (i.e. there was no evidence for changes in binding cooperativity as a result of the mutations studied). Standard errors for the fitted parameters were calculated in GraphPad Prism.

#### Magnetic tweezers

##### Instrument and samples

We used a home-made magnetic tweezers setup similar in design to that described in (58, 59) and detailed in (60). In brief, images of 1 μm superparamagnetic beads tethered to the surface of a glass slide by DNA constructs are acquired with a 100x oil immersion objective and a CCD camera. Real-time image analysis is used to determine the spatial coordinates of beads with nm accuracy in *x*, *y* and *z*. A step-by-step motor located above the sample moves a pair of magnets allowing the application of stretching forces to the bead-DNA system. Applied forces can be quantified from the Brownian excursions of the bead and the extension of the DNA tether. Unless specified otherwise, data were acquired at 120 Hz and filtered down to 3 Hz for representation and analysis

Fabrication of DNA substrates for MT experiments containing a single *parS* sequence with biotins and digoxigenins at the tails was described in (7). The DNA substrates were incubated with 1 μm streptavidin-coated beads (MyOne, Invitrogen) for 10 minutes. Then, the DNA-bead complex was injected in a liquid cell functionalized with anti-digoxigenin antibodies (Roche) and incubated for 10 minutes before applying force. In a first step, visual inspection allows identification and selection of tethered DNA molecules. Torsionally-constrained molecules and beads with more than a single DNA molecule were identified from its distinct rotation-extension curves. Double or multiple tethers were discarded of further analysis in this work. All the experiments were performed in a reaction buffer composed of 100 mM NaCl, 50 mM Tris-HCl or HEPES-KOH pH 7.5, 100 μg/ml BSA and 0.1% Tween 20.

##### CTD-induced decondensation

Once selected single torsionally-relaxed DNA molecules, 1 μM ParB2 was incubated for 2-3 minutes and condensation was induced by decreasing the force to 0.34 pN. Immediately after, one cell-volume of reaction buffer containing 5 μM CTD and 1 μM ParB2 was flown at a constant flow velocity of 16 μl/min. In control experiments where only 1 μM ParB2 was flown, the reaction was supplemented with a volume of storage buffer equal to that used in the CTD experiments and thus maintaining the ionic conditions.

To have a measurement of the degree of induced decondensation, we determined a condensation ratio, Cr (Figure 6D), which was calculated simply as:

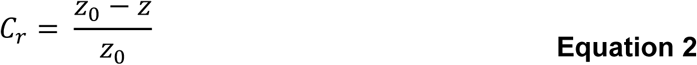
where *z_0_* is the expected extension at 0.34 pN measured before ParB injection and *z* is the equilibrium extension after induced-decondensation. *z* was determined from average extensions of 120 data points at 390 seconds after the cell volume was completely exchanged. These data were acquired at 60 Hz and filtered down to 3 Hz.

##### Force-extension curves and work calculation

Force-extension curves were obtained by decreasing the applied force in steps from 4 pN to ∼ 0.02 pN for a total measuring time of 13 minutes. This procedure is initially performed for bare DNA molecules. Then, the force is reset to 4 pN and ParB variants are flown and incubated for 2 minutes before starting the measurement of a new force-extension curve using the same magnet positions in absence of proteins. In every case, the force applied to each bead was calculated from the force-extension data of bare DNA molecules.

The work done during condensation (Δ*W*) can be calculated by the difference in work between the force-extension curve in the presence of ParB variants and that of bare DNA (Equation 3), where *z_max_* is the extension at the maximum applied force of 4 pN. Integrals were calculated using the trapezoidal rule using OriginLab software.

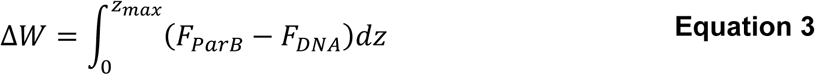

#### Native mass spectrometry

Mass spectra were collected using a Synapt G2 HDMS T-wave ion mobility mass spectrometer (Waters) with nano-electrospray using *in-house* made gold-coated borosilicate capillaries. Protein only experiments required buffer exchange to 300 mM NH_4_Ac (pH 6.9) using Micro Bio-Spin P-6 Gel columns (Bio-Rad). In analysis of CTD-DNA interactions, mixtures of CTD and 15 bp hairpin DNA were co-incubated for 5 mins at 30 °C prior to buffer exchange. The sequence of the 15 bp DNA hairpin was 5’- GCATAGCGTACATCA TTCCC TGATGTACGCTATGC-3’. CTD samples were loaded at 10 or 50 μM. The following parameters were applied to preserve non-covalent interactions (61, 62): backing pressure ca. 5 mbar (adjusted with Speedivalve), source pressure 5.8×10^−3^ mbar, trap pressure 4.4×10^−2^ mbar; capillary voltage 1.3-1.7 kV, sampling cone 20-60 V, extraction cone 1 V, trap and transfer collision energy 10-25 V and 2-5 V, trap DC bias 35-45 V, IMS wave velocity 300-750 m/s, IMS wave height 40 V, helium cell gas flow 180 ml/min, IMS gas flow 90 ml/min (IMS gas cell pressure ca. 3.1 mbar) and source temperature 30 °C. The measured mass of CTD was 8096.1 ± 0.2, which matches well with the calculated value of 8096.1.

#### AUC

Sedimentation velocity experiments were conducted in an Optima XL-A analytical ultracentrifuge using an An-60 Ti rotor (Beckman) at 20 °C. 420 μl volume solutions of 250 μM ParB CTD were prepared in storage buffer with 10% (v/v) glycerol and loaded into a sedimentation velocity cell with sapphire windows and a buffer only reference channel. A rotor speed of 60,000 rpm was employed, with absorbance scans (A_280_) taken across a radial range of 5.85 to 7.25 cm at 2 min intervals to a total of 200 scans. Data were fitted (baseline, meniscus, frictional coefficient (f0), and time- and radial-invariant noise) to a continuous c(s) distribution model using SEDFIT version 9.4, at a 95% confidence level (63, 64). The partial specific volume (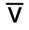) of the CTD and storage buffer density and viscosity were calculated using Sednterp (65). Residuals are shown as a grayscale bitmap where the vertical axis lists each of the 200 scans (with scan 1 at the top) and the horizontal axis depicts radial position over which the data were fitted. Shade indicates variance between fitted and raw data.

#### *In vivo* fluorescence imaging

Nutrient agar (Oxoid) was used for routine selection and maintenance of *B. subtilis* strains. Supplements were added as required: chloramphenicol (5 μg/ml), erythromycin (1 μg/ml), kanamycin (2 μg/ml), spectinomycin (50 μg/ml), tetracycline (10 μg/ml), zeomycin (10 μg/ml), and ampicillin (200 μg/ml). Cells were grown in defined minimal medium base (Spizizen minimal salts supplemented with Fe-NH4-citrate (1 μg/ml), MgSO4 (6 mM), CaCl2 (100 μM), MnSO4 (130 μM), ZnCl2 (1 μM), thiamine (2 μM)) supplemented with casein hydrolysate (200 μg/ml) and/or various carbon sources (succinate (2.0%), glucose (2.0%)). Supplements were added as required: tryptophan (20 μg/ml), erythromycin (1 μg/ml), spectinomycin (50 μg/ml), IPTG (1 mM). Standard techniques were used for strain construction (66). *B. subtilis* competent cells were transformed using an optimised two-step starvation procedure as described (67, 68). All plasmids and strains were verified by sequencing.

To visualise cells, starter cultures were grown at 37 °C overnight in SMM-based medium supplemented with tryptophan (20 μg/ml), casein hydrolysate (200 μg/ml), succinate (2.0%), then diluted 1:100 into fresh medium supplemented with glucose (2.0%) and with/without 1 mM IPTG (as indicated) and allowed to achieve early exponential growth (OD600 0.3-0.4). Cells were mounted on ∼1.2% agar pads (0.25X minimal medium base) and a glass coverslip was placed on top. To visualise individual cells the cell membrane was stained with 0.4 μg/ml FM5-95 (Molecular Probes). Microscopy was performed on an inverted epifluorescence microscope (Nikon Ti) fitted with a Plan-Apochromat objective (Nikon DM 100x/1.40 Oil Ph3). Light was transmitted from a 300 Watt xenon arc-lamp through a liquid light guide (Sutter Instruments) and images were collected using a CoolSnap HQ^2^ cooled CCD camera (Photometrics). All filters were Modified Magnetron ET Sets from Chroma. Digital images were acquired and analysed using METAMORPH software (version V.6.2r6).

#### ChIP-qPCR

To determine the amount of ParB bound to the chromosome by ChIP-qPCR, starter cultures were grown overnight at 30 °C in SMM-based medium supplemented with tryptophan (20 μg/ml), casein hydrolysate (200 μg/ml) and succinate (2.0%), then diluted 1:100 into fresh medium supplemented with glucose (2.0%) and allowed to grow to an A_600_ of 1. Samples were treated with sodium phosphate (final concentration 10 mM) and cross-linked with formaldehyde (final concentration 1%) for 10 mins at room temperature, followed by a further incubation for 30 mins at 4 °C. Cells were pelleted at 15 °C and washed three times with PBS (pH 7.3). Cell pellets were resuspended in 500 μl of lysis buffer (50 mM NaCl, 10 mM Tris-HCl pH 8.0, 20% sucrose, 10 mM EDTA, 100 μg/ml RNase A, ¼ complete mini protease inhibitor tablet (Roche), 2000K u/μl Ready-Lyse lysozyme (Epicentre)) and incubated at 37 °C for 45 mins to degrade the cell wall. 500 μl of IP buffer (300 mM NaCl, 100 mM Tris-HCl pH 7.0, 2% Triton X-100, ¼ complete mini protease inhibitor tablet (Roche), 1 mM EDTA) was added to lyse the cells and the mixture was incubated at 37 °C for a further 10 mins before cooling on ice for 5 mins. To shear DNA to an average size of ∼500 to 1000 bp samples were sonicated (40 amp) four times at 4 °C. The cell debris was removed by centrifugation at 15 °C and the supernatant transferred to a fresh Eppendorf tube. To determine the relative amount of DNA immunoprecipitated compared to the total amount of DNA, 100 μl of supernatant was removed, treated with Pronase (0.5 mg/ml) for 10 mins at 37 °C before SDS (final concentration 0.67%) was added, and stored at 4 °C.

To immunoprecipate protein-DNA complexes, 800 μl of the remaining supernatant was incubated with polyclonal anti-ParB antibodies (Eurogentec) for 1 hr at room temperature. Protein-G Dynabeads (750 μg, Invitrogen) were equilibrated by washing with bead buffer (100 mM Na_3_PO_4_, 0.01% Tween 20), resuspended in 50 μl of bead buffer, and then incubated with the sample supernatant for 3 hrs at room temperature. The immunoprecipated complexes were collected by applying the mixture to a magnet and washed once with the following buffers for 5 mins in the respective order: 0.5X IP buffer; 0.5X IP buffer + NaCl (500 mM); stringent wash buffer (250 mM LiCl, 10 mM Tris-HCl pH 8.0, 0.5% Tergitol-type NP-40, 0.5% C_24_H_39_NaO_4_, 10 mM EDTA). Finally, the complexes were washed a further three times with TET buffer (10 mM Tris-HCl pH 8.0, 1 mM EDTA, 0.01% Tween 20) and resuspended in 112 μl of TE buffer (10 mM Tris-HCl pH 8.0, 1 mM EDTA). Formaldehyde crosslinks of both the total DNA and the immunoprecipate was reversed by incubation at 65 °C for 16 hrs. The reversed DNA was then removed from the magnetic beads and transferred to a clean PCR tube and stored at 4 °C for qPCR analysis.

To measure the amount of DNA bound to ParB, GoTaq (Promega) qPCR mix was used for the PCR reactions and qPCR was performed in a Rotor-Gene Q Instrument (Qiagen) using serial dilution of the immunoprecipitate and the total DNA control as the template. Oligonucleotide primers were then designed that amplify at an interval of ∼500-1000 bp away from *parS*^359°^ and were typically 20-25 bases in length and amplified a ∼200-300 bp PCR product (Table S2).

#### Western blot analysis

Proteins of the whole cell extract were separated by electrophoresis using a NuPAGE 4-12% Bis-Tris gradient gel in MES buffer (Life Technologies) and transferred to a Hybond-P PVDF membrane (GE Healthcare Life Sciences) using a semi dry apparatus (Hoefer Scientific Instruments). Polyclonal primary antibodies were used to probe protein of interest and then detected with an anti-rabbit horseradish peroxidase-linked secondary antibody using an ImageQuant LAS 4000 mini digital imaging system (GE Healthcare Life Sciences).

---

**Figure S1.**
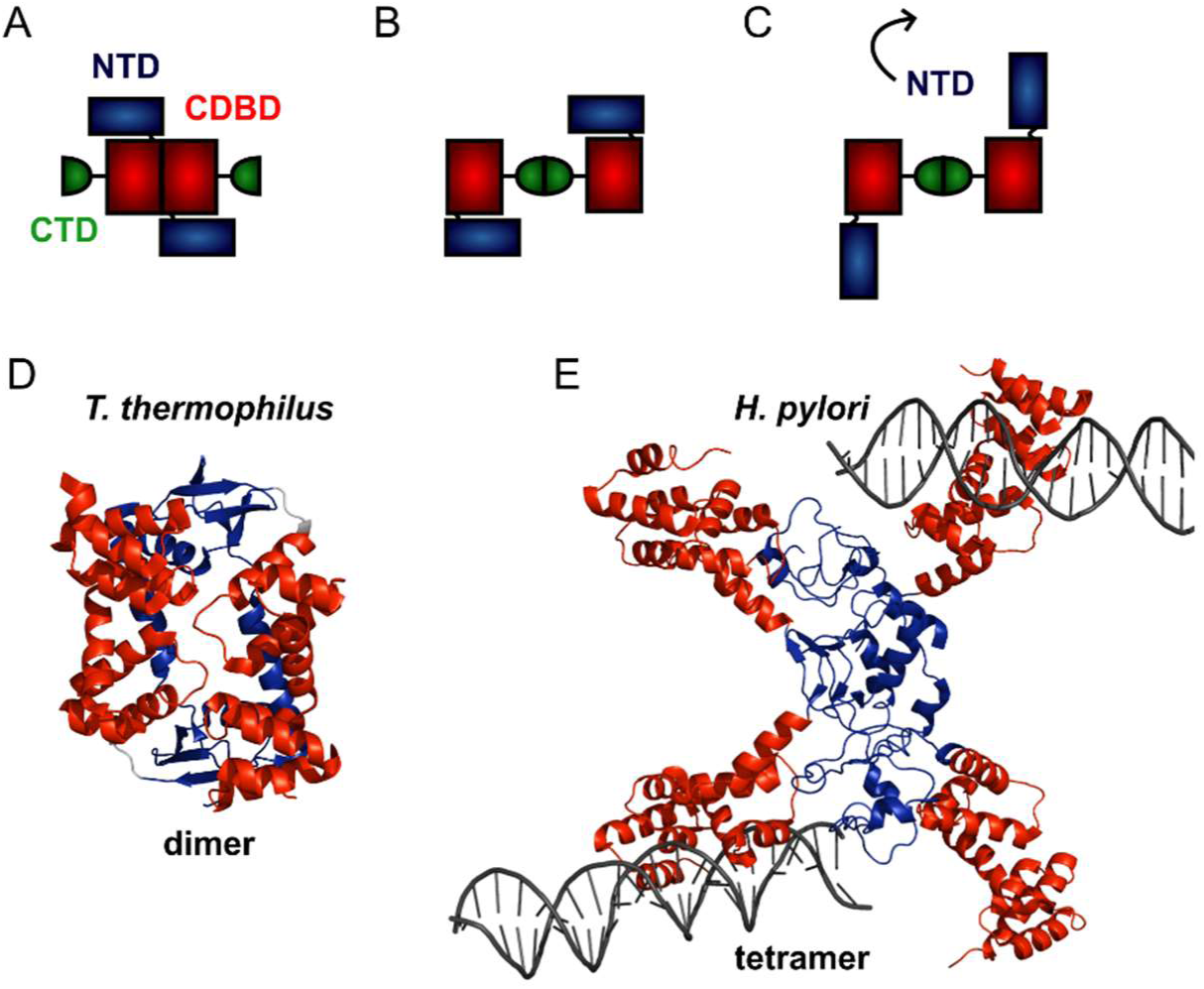
Structural models for genomic ParB. (A-C) Schematics of different domain arrangements for dimeric *B. subtilis* ParB. Whilst it is known that ParB is dimeric in solution, the primary dimerisation interface is unclear and possibly condition-dependent. Structural evidence suggests that this could occur between the N-terminal and central DNA-binding domains or between the C-terminal domains. Additional structural evidence suggests that the NTD may flip away from the CDBD to form higher order ParB oligomers via a tetrameric interface. (D) Crystal structure of a dimer of the NTD and CDBD of *T. thermophilus* ParB (PDB entry IVZ0) (14). (E) Crystal structure of a tetramer of the NTD and CDBD of *H. pylori* ParB bound to 2 *parS*-containing 24 bp dsDNA (PDB entry 4MUK) (16). The structures are colour-coded as in the primary structure diagram (Figure 1, main text).

**Figure S2.**
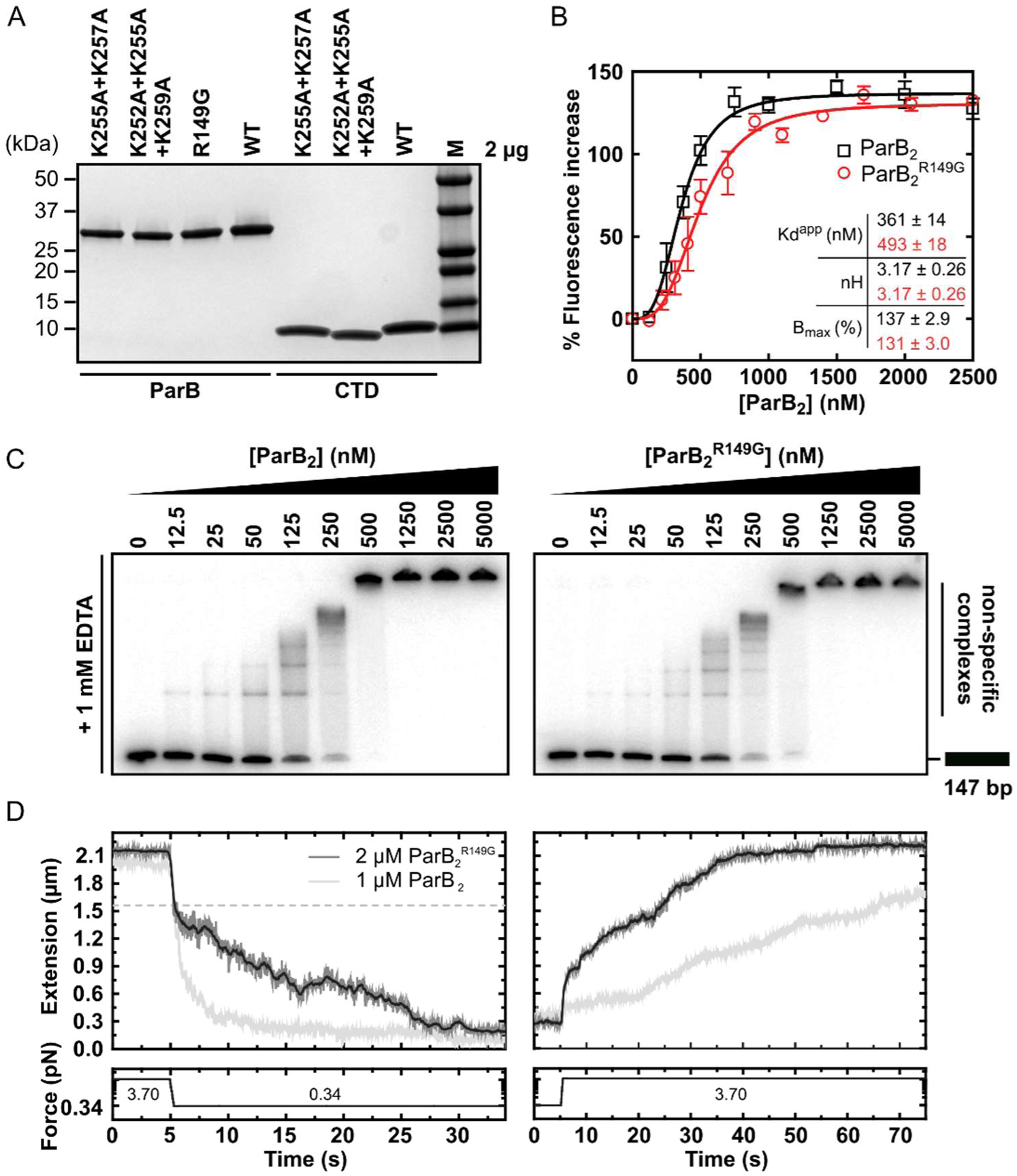
The helix-turn-helix motif is dispensable for non-specific binding and condensation. (A) SDS-PAGE of wild type ParB and mutant protein preparations. (B) Non-specific DNA binding isotherms for wild type ParB and ParB^R149G^ measured using a PIFE assay. The data were fitted to the Hill equation. Error bars represent the standard errors from 3 independent experiments. Standard errors of fitted parameters were calculated in GraphPad Prism. (C) TBE-EMSAs for wild type ParB and ParB^R149G^ binding 147 bp DNA. (D) Left panel: representative condensation traces for wild type ParB and ParB^R149G^, monitored by magnetic tweezers. At a 3.70 pN stretching force, ParB was introduced to single DNA tethers. Following incubation, force was reduced to 0.34 pN to induce condensation. Expected DNA extension under 0.34 pN and in the absence of ParB is indicated by the dashed line. Right panel: representative force-induced decondensation traces for wild type ParB and ParB^R149G^.

**Figure S3.**
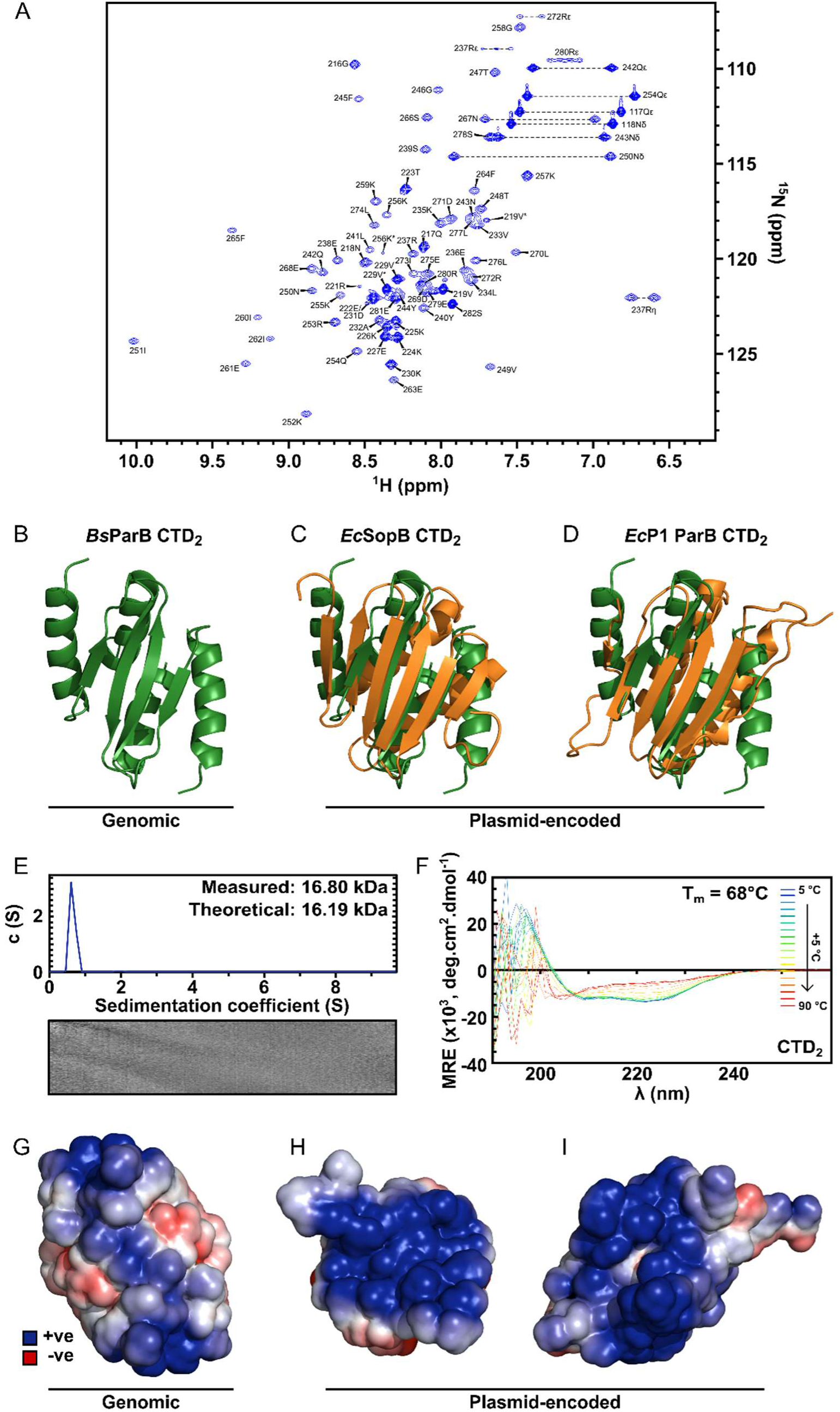
Solution NMR structure of the dimeric ParB CTD domain. (A) Assigned ^1^H-^15^N HSQC spectra of CTD. Arginine ε and η peaks are aliased once and twice, respectively. Starred peaks indicate additional (usually minor) species. (B) The C-terminal domain of *B. subtilis* ParB (green). (C) Alignment of the secondary structure elements of the CTDs of *B. subtilis* ParB and *E. coli* plasmid-encoded SopB (orange) (32). PDB entry: 3K75. (D) Alignment of the secondary structure elements of the CTDs of *B. subtilis* ParB and *E. coli* plasmid-encoded P1 ParB (orange) (69). PDB entry: 2NTZ. (E) ParB sedimentation velocity c(s) distribution fit at 20°C. Residuals are shown as a grayscale bitmap where the vertical axis lists each scan and the horizontal axis depicts radial position over which data were fitted. Shade indicates variance between fitted and raw data. RMSD = 0.0079. (F) CD spectra over a thermal denaturation scan (5 to 90 °C) for 10 μM CTD**2** in PBS. (G-I) Continuum electrostatics calculations were prepared using the PDB2PQR web server (41) and the APBS plugin for PyMOL (42). G to I are *B. subtilis* ParB, *E. coli* F-plasmid SopB and *E. coli* P1 ParB, respectively.

**Figure S4.**
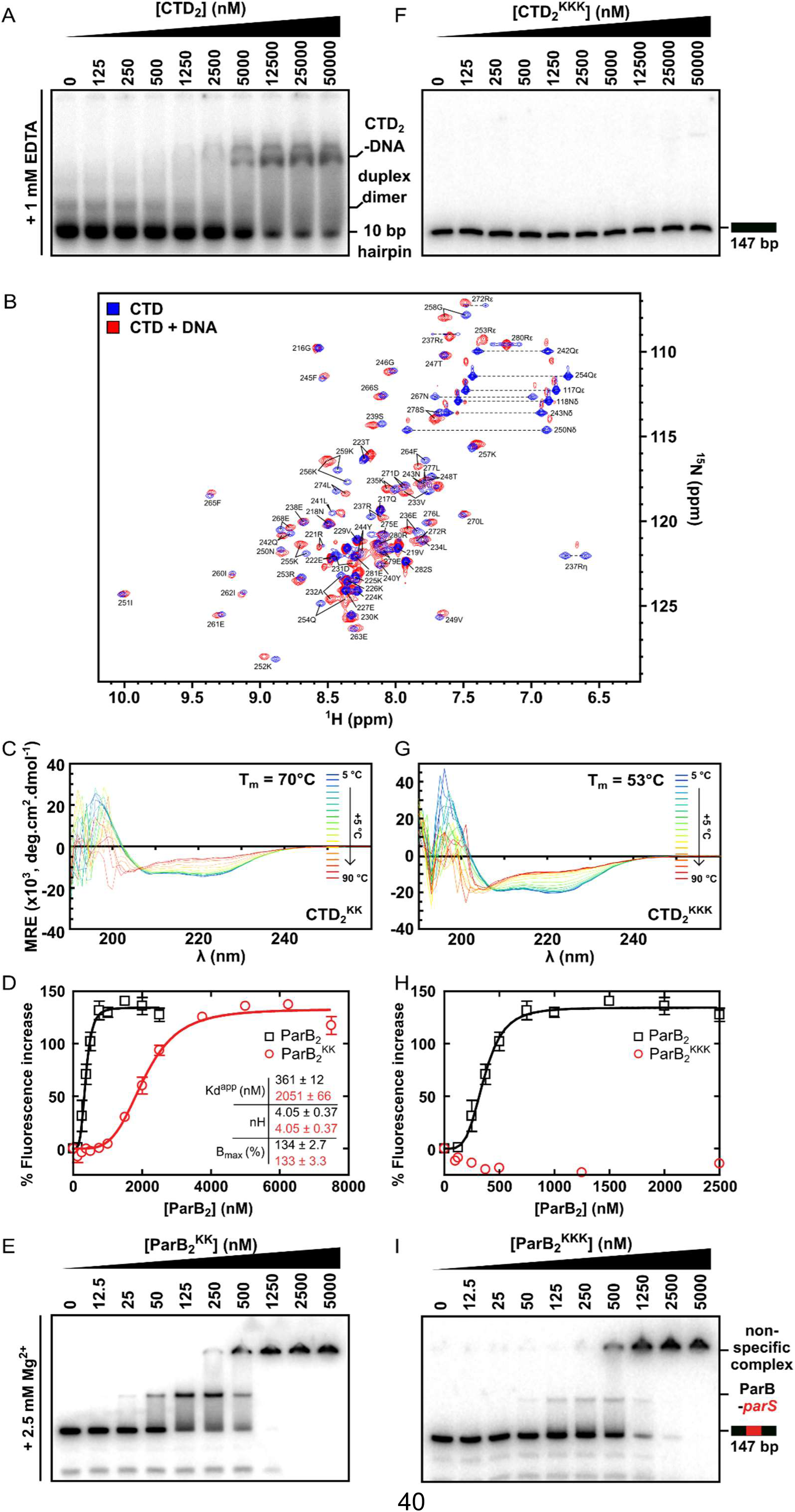
The CTD binds DNA via a lysine-rich surface. (A) TBE-EMSA showing binding of the CTD to the 10 bp hairpin DNA substrate used for NMR experiments. (B) Assigned ^1^H-^15^N HSQC spectra of the CTD prior (blue) and after (red) titration with a 10 bp hairpin DNA substrate. Arginine ε and η peaks are aliased once and twice, respectively. (C) CD spectra over a thermal denaturation scan (5 to 90 °C) for 10 μM CTD**2**^KK^ in PBS. (D) PIFE assay showing binding of full length ParB^KK^ to non-specific DNA. The data were fitted to the Hill equation. Error bars represent the standard errors from 3 independent experiments. Standard errors of fitted parameters were calculated in GraphPad Prism. (E) TBM-EMSA showing binding of full length ParB^KK^ to *parS*-containing DNA. ParB^KK^ is able to form specific complexes. (F) TBE-EMSA showing no binding of CTD^KKK^ to DNA at concentrations as high as 50 μM dimer. (G) CD spectra for 10 μM CTD**2**^KKK^ recorded as in C. (H) As in D, using ParB^KKK^. H uses the same wild-type ParB dataset as D. (I) TBM-EMSA showing binding of full length ParB^KKK^ to *parS*-containing DNA. This mutant protein is also able to form specific complexes.

**Figure S5.**
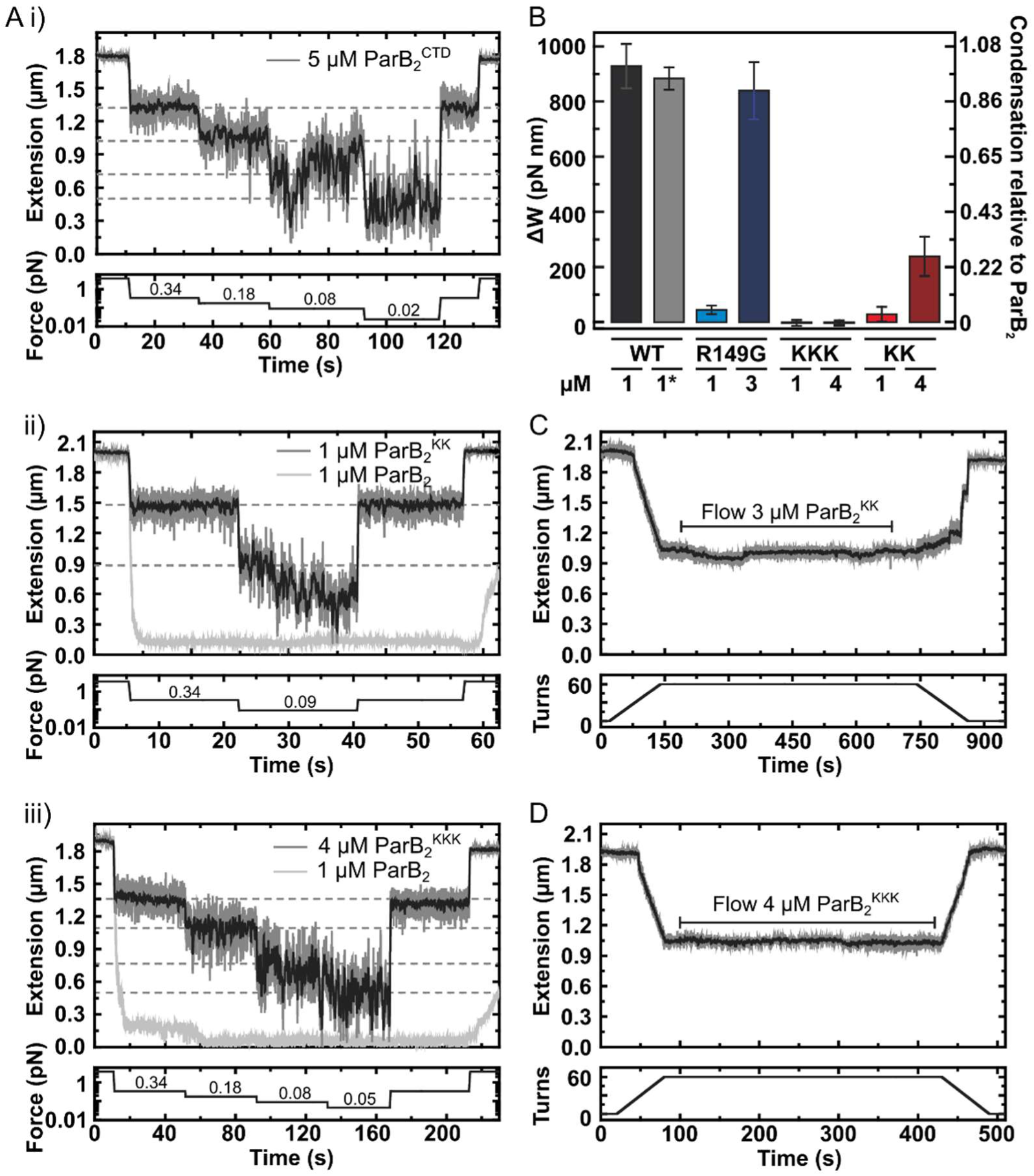
DNA binding by the CTD is required for efficient DNA condensation *in vitro*. (A) Representative traces of condensation assays monitored by magnetic tweezers. The expected DNA extension in the absence of ParB at each force is indicated by the dashed lines. i), ii) and ii) use CTD, ParB^KK^ and ParB^KKK^, respectively. (B) A comparison of the work produced by full length ParB variants in condensing DNA. Errors are standard error of the mean. * indicates this experiment used nsDNA rather than *parS*-containing. (C) Representative trace of a ParB^KK^ plectoneme stabilisation assay showing hysteresis of DNA extension upon DNA rewinding. (D) Representative trace of a ParB^KKK^ plectoneme stabilisation assay showing no hysteresis upon DNA rewinding.

**Figure S6.**
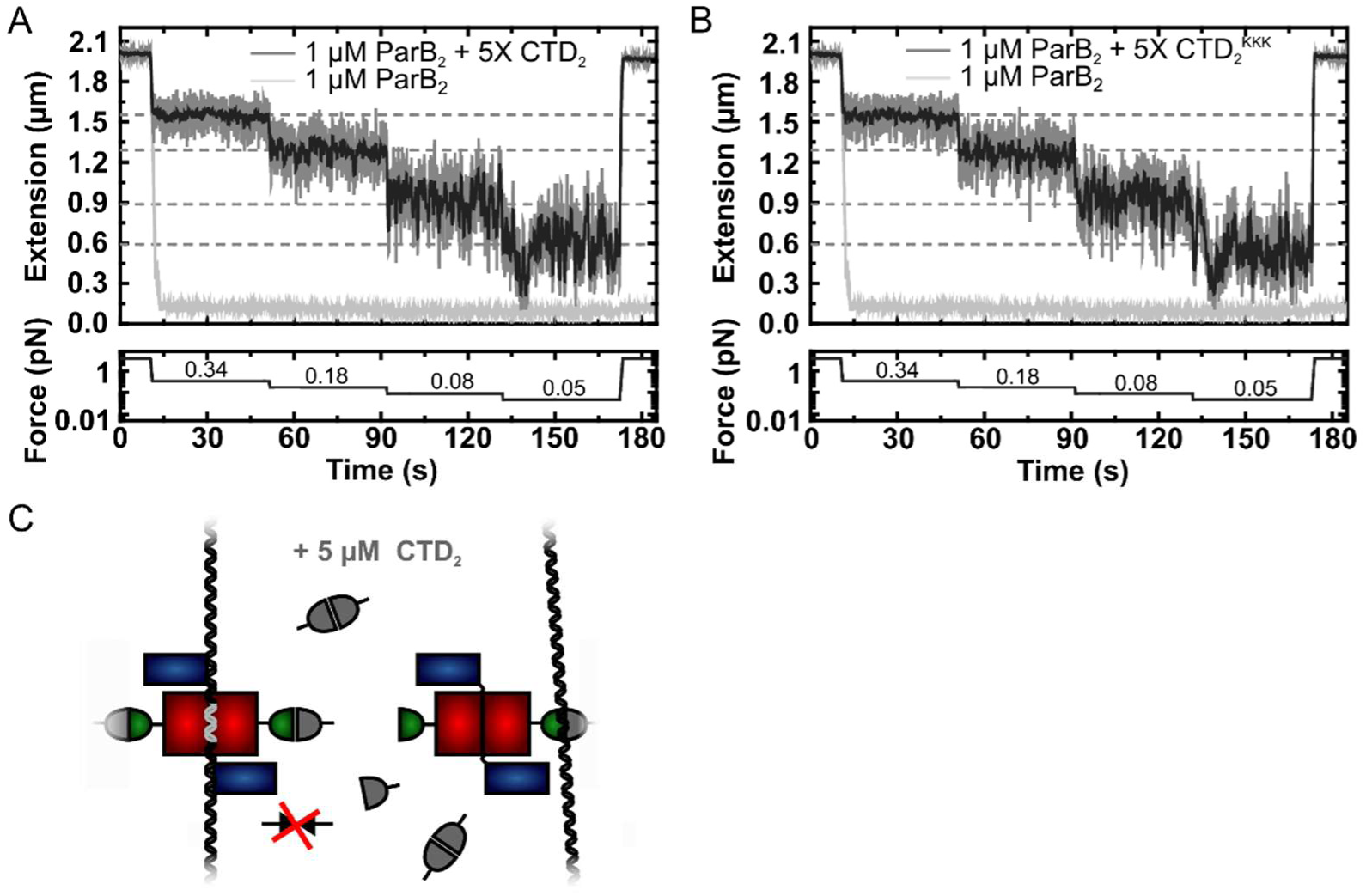
The CTD of ParB both inhibits and disrupts ParB-dependent DNA condensation. (A) Representative traces of condensation assays for ParB in the presence and absence of excess free CTD. The expected DNA extension in the absence of ParB at each force is indicated by the dashed lines. (B) As in A, but using the CTD^KKK^ variant in place of the CTD. (C) Model for how the addition of an excess of the CTD can inhibit ParB-DNA networks. DNA-binding deficient CTD variants are equally efficient competitors, indicating that interference with the CTD dimerisation/bridging interface is important for the inhibitory effect.

**Figure S7.**
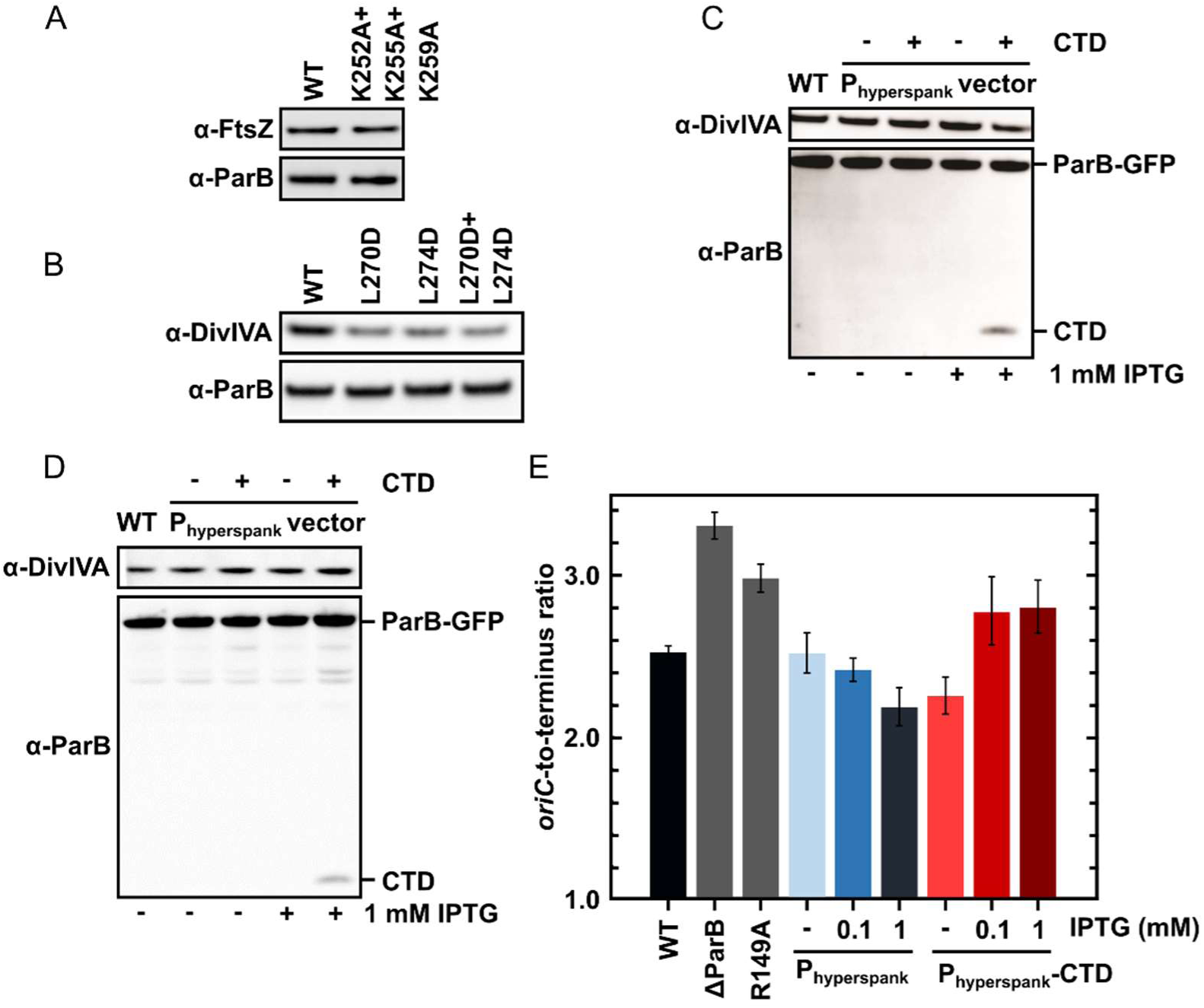
DNA-binding and dimerisation by the CTD is critical for ParB function *in vivo*. (A-B) Western blot analysis. Expression levels of parB^L270D+L274D^ and parB^K252A+K255A+K259A^ with a C-terminal gfp fusion were similar to wild type ParB. DivIVA is an abundant cell division protein that was used as a loading control. (C) Wild type ParB-GFP levels of expression were unaffected by overexpression of CTD. (D) As in C using the construct for ChIP-qPCR (Figure 7D). (E) Marker frequency analysis was used to measure the rate of DNA replication initiation. Cells were grown in slow growth media and allowed to achieve at least three mass doublings (A_600_ 0.3 to 0.5) before DNA was extracted. Regulation of DnaA by Soj was affected by overexpression of CTD. R149A is a *parS* DNA-binding mutant.

**Table S1.** NMR assignment, structure calculation and validation statistics. ^a^commonly assigned groups (i.e. excluding OH, Asp/Glu side-chain carbonyl, Lys amide and Arg guanidinium groups as well as tertiary aromatic carbons); ^b^residues 229-282; ^c^values reported by ARIA 2.3 (51); ^d^ordered residues (230-254, 259-278) as calculated by PSVS 1.5 (56); ^e^values reported by Procheck (72); ^f^value reported by PDB validation software; ^g^ residues in secondary structure (231-245, 249-254, 257-264, 267-277).

|  |  |
| --- | --- |
| **Degree of assignment<sup>a,b</sup>** |  |
| Backbone (Cα, C', N and H<sup>N</sup>) (%) | 98.5 |
| Side-chain H (%) | 97.5 |
| Side-chain non-H (%) | 96.2 |
| **Number of restraints (per monomer)** |  |
| NOE restraints |  |
| Intra-residue ($\vert i-j\vert = 0$) | 477 |
| Sequential ($\vert i-j\vert = 1$) | 319 |
| Medium range ($2 \leq \vert i-j\vert < 5$) | 324 |
| Long range ($\vert i-j\vert \geq 5$) | 392 |
| Ambiguous | 481 |
| Total | 1993 |
| H-bond restraints (intra-/intermonomer) | 24 / 4 |
| Dihedral angle restraints ($\Phi/\Psi$) | 40 / 40 |
| **Restraint statistics<sup>c</sup>** |  |
| r.m.s. of NOE violations (Å) | 0.09 ± 0.01 |
| r.m.s. of H-bond violations (Å) | 0.19 ± 0.11 |
| r.m.s. of dihedral violations (°) | 0.15 ± 0.07 |
| **r.m.s. from idealised covalent geometry<sup>c</sup>** |  |
| Bonds (Å) | 0.0032 ± 0.0001 |
| Angles (°) | 0.48 ± 0.015 |
| Impropers (°) | 1.20 ± 0.12 |
| **Structural quality** |  |
| Ramachandran statistics<sup>e,b,d</sup> |  |
| Most favoured regions (%) | 87.4 / 93.6 |
| Allowed regions (%) | 10.8 / 6.4 |
| Generously allowed regions (%) | 0.4 / 0.0 |
| Disallowed regions (%) | 1.4 / 0.0 |
| CING (57) % ROG scores (R/O/G)<sup>b</sup> | 19 / 15 / 67 |
| Verify3D (70) Z-score<sup>b</sup> | -0.16 |
| Prosa II (71) Z-score<sup>b</sup> | -1.78 |
| Procheck (72) Z-score ($\Phi/\Psi$)<sup>b,d</sup> | 0.79 |
| Procheck (72) Z-score (all)<sup>b,d</sup> | -0.35 |
| MolProbity (73) Z-score<sup>b</sup> | -2.51 |
| No. of close contacts<sup>b,f</sup> | 4 |
| **Coordinates precision (rmsd)** <sup>b/d/g</sup> |  |
| All backbone atoms (Å) | 0.8 / 0.5 / 0.4 |
| All heavy atoms (Å) | 1.0 / 0.8 / 0.7 |

**Table S2.**
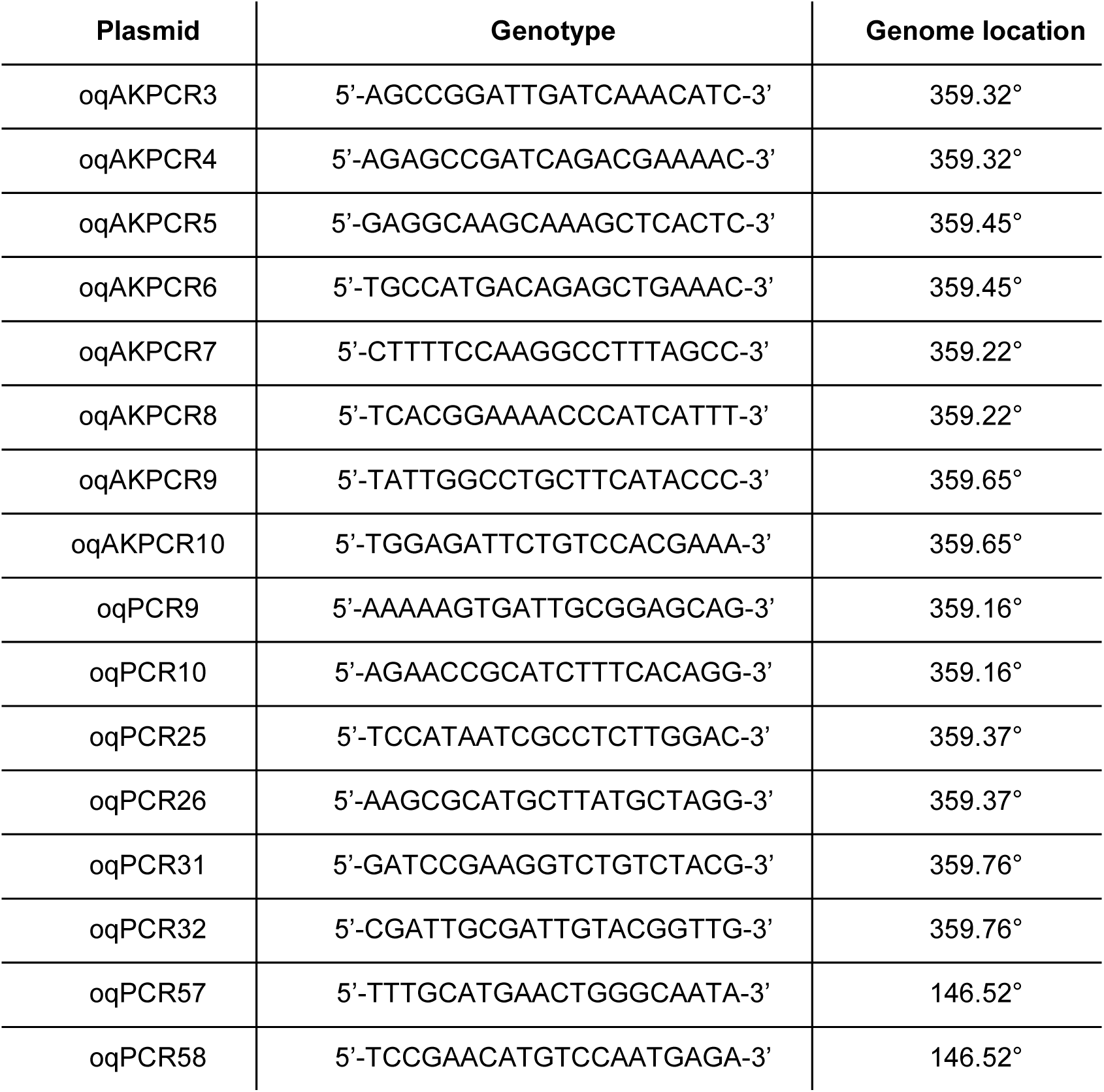
Primer sequences used in ChIP-qPCR

## References

1 Wang, X., Tang, O.W., Riley, E.P. and Rudner, D.Z. (2014) The SMC Condensin Complex Is Required for Origin Segregation in Bacillus subtilis. Curr. Biol., 10.1016/j.cub.2013.11.050.

2 Breier, A.M. and Grossman, A.D. (2007) Whole-genome analysis of the chromosome partitioning and sporulation protein Spo0J (ParB) reveals spreading and origin-distal sites on the Bacillus subtilis chromosome. Mol. Microbiol., 64, 703–18.

3 Murray, H., Ferreira, H. and Errington, J. (2006) The bacterial chromosome segregation protein Spo0J spreads along DNA from parS nucleation sites. Mol. Microbiol., 61, 1352–61.

4 Lynch, a S. and Wang, J.C. (1995) SopB protein-mediated silencing of genes linked to the sopC locus of Escherichia coli F plasmid. Proc. Natl. Acad. Sci. U. S. A., 92, 1896–900.

5 Rodionov, O., Lobocka, M. and Yarmolinsky, M. (1999) Silencing of Genes Flanking the P1 Plasmid Centromere. Science (80-.)., 283, 546–549.

6 Graham, T.G.W., Wang, X., Song, D., Etson, C.M., van Oijen, A.M., Rudner, D.Z. and Loparo, J.J. (2014) ParB spreading requires DNA bridging. Genes Dev., 28, 1228–38.

7 Taylor, J.A., Pastrana, C.L., Butterer, A., Pernstich, C., Gwynn, E.J., Sobott, F., Moreno-Herrero, F. and Dillingham, M.S. (2015) Specific and non-specific interactions of ParB with DNA: implications for chromosome segregation. Nucleic Acids Res., 10.1093/nar/gku1295.

8 Broedersz, C.P., Wang, X., Meir, Y., Loparo, J.J., Rudner, D.Z. and Wingreen, N.S. (2014) Condensation and localization of the partitioning protein ParB on the bacterial chromosome. Proc. Natl. Acad. Sci., 111, 8809–14.

9 Sanchez, A., Cattoni, D.I., Walter, J.-C., Rech, J., Parmeggiani, A., Nollmann, M. and Bouet, J.-Y. (2015) Stochastic Self-Assembly of ParB Proteins Builds the Bacterial DNA Segregation Apparatus. Cell Syst., 1, 163–173.

10 Bouet, J.-Y. and Funnell, B.E. (1999) P1 ParA interacts with the P1 partition complex at parS and an ATP-ADP switch controls ParA activities. EMBO J., 18, 1415–24.

11 Davey, M.J. and Funnell, B.E. (1997) Modulation of the P1 plasmid partition protein ParA by ATP, ADP, and P1 ParB. J. Biol. Chem., 272, 15286–15292.

12 Davis MA, Martin KA, A.S. (1992) Biochemical activities of the parA partition protein of the P1 plasmid. Mol. Microbiol., 6, 1141–7.

13 Radnedge, L., Youngren, B., Davis, M. and Austin, S. (1998) Probing the structure of complex macromolecular interactions by homolog specificity scanning: The P1 and P7 plasmid partition systems. EMBO J., 17, 6076–6085.

14 Leonard, T.A., Butler, P.J.G. and Löwe, J. (2004) Structural analysis of the chromosome segregation protein Spo0J from Thermus thermophilus. Mol. Microbiol., 53, 419–32.

15 Schumacher, M.A. and Funnell, B.E. (2005) Structures of ParB bound to DNA reveal mechanism of partition complex formation. Nature, 438, 516–9.

16 Chen, B.-W., Lin, M.-H., Chu, C.-H., Hsu, C.-E. and Sun, Y.-J. (2015) Insights into ParB spreading from the complex structure of Spo0J and parS. Proc. Natl. Acad. Sci. U. S. A., 112, 6613–8.

17 Schumacher, M.A., Mansoor, A. and Funnell, B.E. (2007) Structure of a four-way bridged ParB-DNA complex provides insight into P1 segrosome assembly. J. Biol. Chem., 282, 10456–64.

18 Autret, S., Nair, R. and Errington, J. (2001) Genetic analysis of the chromosome segregation protein Spo0J of Bacillus subtilis: evidence for separate domains involved in DNA binding and interactions with Soj protein. Mol. Microbiol., 41, 743–55.

19 Gruber, S. and Errington, J. (2009) Recruitment of condensin to replication origin regions by ParB/SpoOJ promotes chromosome segregation in B. subtilis. Cell, 137, 685–96.

20 Lin, D.C. and Grossman, a D. (1998) Identification and characterization of a bacterial chromosome partitioning site. Cell, 92, 675–85.

21 Real, G., Autret, S., Harry, E.J., Errington, J. and Henriques, A.O. (2005) Cell division protein DivIB influences the Spo0J/Soj system of chromosome segregation in Bacillus subtilis. Mol. Microbiol., 55, 349–67.

22 Glaser, P., Sharpe, M.E., Raether, B., Perego, M., Ohlsen, K. and Errington, J. (1997) Dynamic, mitotic-like behavior of a bacterial protein required for accurate chromosome partitioning. Genes Dev., 11, 1160–1168.

23 Lin, D.C., Levin, P. a and Grossman, A.D. (1997) Bipolar localization of a chromosome partition protein in Bacillus subtilis. Proc. Natl. Acad. Sci. U. S. A., 94, 4721–6.

24 Lewis, P.J. and Errington, J. (1997) Direct evidence for active segregation of oriC regions of the Bacillus subtilis chromosome and co-localization with the SpoOJ partitioning protein. Mol. Microbiol., 25, 945–954.

25 Marston, a L. and Errington, J. (1999) Dynamic movement of the ParA-like Soj protein of B. subtilis and its dual role in nucleoid organization and developmental regulation. Mol. Cell, 4, 673–682.

26 Murray, H. and Errington, J. (2008) Dynamic Control of the DNA Replication Initiation Protein DnaA by Soj/ParA. Cell, 135, 74–84.

27 Bingle, L.E.H., Macartney, D.P., Fantozzi, A., Manzoor, S.E., Thomas, C.M. and Karn, J. (2005) Flexibility in repression and cooperativity by KorB of broad host range IncP-1 plasmid RK2. J. Mol. Biol., 349, 302–316.

28 Wang, X., Brandão, H.B., Le, T.B.K., Laub, M.T. and Rudner, D.Z. (2017) Bacillus subtilis SMC complexes juxtapose chromosome arms as they travel from origin to terminus. Science (80-.)., 355, 524–527.

29 Gerdes, K., Møller-Jensen, J. and Bugge Jensen, R. (2000) Plasmid and chromosome partitioning: surprises from phylogeny. Mol. Microbiol., 37, 455–66.

30 Schumacher, M. a (2008) Structural biology of plasmid partition: uncovering the molecular mechanisms of DNA segregation. Biochem. J., 412, 1–18.

31 Sanchez, A., Rech, J., Gasc, C. and Bouet, J.-Y. (2013) Insight into centromere-binding properties of ParB proteins: a secondary binding motif is essential for bacterial genome maintenance. Nucleic Acids Res., 41, 3094–103.

32 Schumacher, M.A., Piro, K.M. and Xu, W. (2010) Insight into F plasmid DNA segregation revealed by structures of SopB and SopB-DNA complexes. Nucleic Acids Res., 38, 4514–26.

33 Kusukawa, N., Mori, H., Kondo, A. and Hiraga, S. (1987) Partitioning of the F plasmid: Overproduction of an essential protein for partition inhibits plasmid maintenance. MGG Mol. Gen. Genet., 208, 365–372.

34 Bignell, C. and Thomas, C.M. (2001) The bacterial ParA-ParB partitioning proteins. J. Biotechnol., 91, 1–34.

35 Theophilus, B.D.M. and Thomas, C.M. (1987) Nucleotide sequence of the transcriptional repressor gene korB which plays a key role in regulation of the copy number of broad host range plasmid RK2. Nucleic Acids Res., 15, 7443–7450.

36 Lobocka, M. and Yarmolinsky, M. (1996) P1 plasmid partition: a mutational analysis of ParB. J. Mol. Biol., 259, 366–82.

37 Erdmann, N., Petroff, T. and Funnell, B.E. (1999) Intracellular localization of P1 ParB protein depends on ParA and parS. Proc. Natl. Acad. Sci. U. S. A., 96, 14905–10.

38 Sullivan, N.L., Marquis, K. a and Rudner, D.Z. (2009) Recruitment of SMC by ParB-parS organizes the origin region and promotes efficient chromosome segregation. Cell, 137, 697–707.

39 Bartosik, A.A., Lasocki, K., Mierzejewska, J., Thomas, C.M. and Jagura-Burdzy, G. (2004) ParB of Pseudomonas aeruginosa: interactions with its partner ParA and its target parS and specific effects on bacterial growth. J. Bacteriol., 186, 6983–98.

40 Kusiak, M., Gapczynska, A., Plochocka, D., Thomas, C.M. and Jagura-Burdzy, G. (2011) Binding and spreading of ParB on DNA determine its biological function in Pseudomonas aeruginosa. J. Bacteriol., 193, 3342–55.

41 Dolinsky, T.J., Nielsen, J.E., McCammon, J.A. and Baker, N.A. (2004) PDB2PQR: An automated pipeline for the setup of Poisson-Boltzmann electrostatics calculations. Nucleic Acids Res., 32, 665–667.

42 Carlson, M.L. and H. (2006) MG Lerner and HA Carlson. APBS plugin for PyMOL.

43 Baker, N.A., Sept, D., Joseph, S., Holst, M.J. and McCammon, J.A. (2001) Electrostatics of nanosystems: application to microtubules and the ribosome. Proc. Natl. Acad. Sci. U. S. A., 98, 10037–41.

44 Goldenberg, O., Erez, E., Nimrod, G. and Ben-Tal, N. (2009) The ConSurf-DB: pre-calculated evolutionary conservation profiles of protein structures. Nucleic Acids Res., 37, D323–D327.

45 Celniker, G., Nimrod, G., Ashkenazy, H., Glaser, F., Martz, E., Mayrose, I., Pupko, T. and Ben-Tal, N. (2013) ConSurf: Using Evolutionary Data to Raise Testable Hypotheses about Protein Function. Isr. J. Chem., 53, 199–206.

46 Williams, C., Rezgui, D., Prince, S.N., Zaccheo, O.J., Foulstone, E.J., Forbes, B.E., Norton, R.S., Crosby, J., Hassan, A.B. and Crump, M.P. (2007) Structural Insights into the Interaction of Insulin-like Growth Factor 2 with IGF2R Domain 11. Structure, 15, 1065–1078.

47 Zwahlen, C., Legault, P., Vincent, S.J.F., Greenblatt, J., Konrat, R. and Kay, L.E. (1997) Methods for measurement of intermolecular NOEs by multinuclear NMR spectroscopy: Application to a bacteriophage λ N-peptide/boxB RNA complex. J. Am. Chem. Soc., 119, 6711–6721.

48 Delaglio, F., Grzesiek, S., Vuister, G., Zhu, G., Pfeifer, J. and Bax, A. (1995) NMRPipe: A multidimensional spectral processing system based on UNIX pipes. J. Biomol. NMR, 6, 277–293.

49 Vranken, W.F., Boucher, W., Stevens, T.J., Fogh, R.H., Pajon, A., Llinas, M., Ulrich, E.L., Markley, J.L., Ionides, J. and Laue, E.D. (2005) The CCPN data model for NMR spectroscopy: Development of a software pipeline. Proteins Struct. Funct. Genet., 59, 687–696.

50 Shen, Y., Delaglio, F., Cornilescu, G. and Bax, A. (2009) TALOS+: A hybrid method for predicting protein backbone torsion angles from NMR chemical shifts. J. Biomol. NMR, 44, 213–223.

51 Rieping, W., Bardiaux, B., Bernard, A., Malliavin, T.E. and Nilges, M. (2007) ARIA2: Automated NOE assignment and data integration in NMR structure calculation. Bioinformatics, 23, 381–382.

52 Linge, J.P., Habeck, M., Rieping, W. and Nilges, M. (2004) Correction of spin diffusion during iterative automated NOE assignment. J. Magn. Reson., 167, 334–342.

53 Bardiaux, B., Bernard, A., Rieping, W., Habeck, M., Malliavin, T.E. and Nilges, M. (2009) Influence of different assignment conditions on the determination of symmetric homodimeric structures with ARIA. Proteins Struct. Funct. Bioinforma., 75, 569–585.

54 Nilges, M., Bernard, A., Bardiaux, B., Malliavin, T., Habeck, M. and Rieping, W. (2008) Accurate NMR Structures Through Minimization of an Extended Hybrid Energy. Structure, 16, 1305–1312.

55 Mareuil, F., Malliavin, T.E., Nilges, M. and Bardiaux, B. (2015) Improved reliability, accuracy and quality in automated NMR structure calculation with ARIA. J. Biomol. NMR, 62, 425–438.

56 Bhattacharya, A., Tejero, R. and Montelione, G.T. (2007) Evaluating Protein Structures Determined by Structural Genomics Consortia. Proteins, 66, 778–795.

57 Doreleijers J.F., Da Silva, A.W.S., Krieger, E., Nabuurs, S.B., Spronk, C.A.E.M., Stevens, T.J., Vranken, W.F., Vriend, G. and Vuister, G.W. (2012) CING: An integrated residue-based structure validation program suite. J. Biomol. NMR, 54, 267–283.

58 Strick, T.R., Allemand, J.F., Bensimon, D. and Croquette, V. (1998) Behavior of supercoiled DNA. Biophys. J., 74, 2016–28.

59 Carrasco, C., Gilhooly, N.S., Dillingham, M.S. and Moreno-Herrero, F. (2013) On the mechanism of recombination hotspot scanning during double-stranded DNA break resection. Proc. Natl. Acad. Sci. U. S. A., 110, E2562–71.

60 Pastrana, C.L., Carrasco, C., Akhtar, P., Leuba, S.H., Khan, S.A. and Moreno-Herrero, F. (2016) Force and twist dependence of RepC nicking activity on torsionally-constrained DNA molecules. Nucleic Acids Res., 44, 8885–8896.

61 Sobott, F., McCammon, M.G., Hernández, H. and Robinson, C. V (2005) The flight of macromolecular complexes in a mass spectrometer. Philos. Trans. A. Math. Phys. Eng. Sci., 363, 379-389-391.

62 Konijnenberg, A., Butterer, A. and Sobott, F. (2013) Native ion mobility-mass spectrometry and related methods in structural biology. Biochim. Biophys. Acta - Proteins Proteomics, 1834, 1239–1256.

63 Schuck, P. (2000) Size-distribution analysis of macromolecules by sedimentation velocity ultracentrifugation and lamm equation modeling. Biophys. J., 78, 1606–19.

64 Brown, P.H. and Schuck, P. (2006) Macromolecular size-and-shape distributions by sedimentation velocity analytical ultracentrifugation. Biophys. J., 90, 4651–61.

65 Hayes, D., Laue, T. and Philo, J. (1995) Program Sednterp: sedimentation interpretation program.

66 Harwood, C.R., and Cutting, S.M. (1990) Molecular Biological Methods for Bacillus. Wiley, New York.

67 Anagnostopoulos, C. and Spizizen, J. (1961) Requirements for Transformation in Bacillus Subtilis. J. Bacteriol., 81, 741–6.

68 Hamoen, L.W., Smits, W.K., de Jong, A., Holsappel, S. and Kuipers, O.P. (2002) Improving the predictive value of the competence transcription factor (ComK) binding site in Bacillus subtilis using a genomic approach. Nucleic Acids Res., 30, 5517–28.

69 Vecchiarelli, A.G., Schumacher, M. a and Funnell, B.E. (2007) P1 partition complex assembly involves several modes of protein-DNA recognition. J. Biol. Chem., 282, 10944–52.

70 Lüthy, R., Bowie, J.U. and Eisenberg, D. (1992) Assessment of protein models with three-dimensional profiles. Nature, 356, 83–5.

71 Sippl, M.J. (1993) Recognition of Errors in 3-Dimensional Structures of Proteins. Proteins-Structure Funct. Genet., 17, 355–362.

72 Laskowski, R. a., MacArthur, M.W., Moss, D.S. and Thornton, J.M. (1993) PROCHECK: a program to check the stereochemical quality of protein structures. J. Appl. Crystallogr., 26, 283–291.

73 Lovell, S.C., Davis, I.W., Adrendall, W.B., de Bakker, P.I.W., Word, J.M., Prisant, M.G., Richardson, J.S. and Richardson, D.C. (2003) Structure validation by C alpha geometry: phi, psi and C beta deviation. Proteins-Structure Funct. Genet., 50, 437–450.

